# The genetic basis of *Drosophila melanogaster* defense against *Beauveria bassiana* explored through evolve and resequence and quantitative trait locus mapping

**DOI:** 10.1101/2021.03.31.437898

**Authors:** Parvin Shahrestani, Elizabeth King, Reza Ramezan, Mark Phillips, Melissa Riddle, Marisa Thornburg, Zachary Greenspan, Yonathan Estrella, Kelly Garcia, Pratik Chowdhury, Glen Malarat, Ming Zhu, Susan M. Rottshaefer, Stephen Wraight, Michael Griggs, John Vandenberg, Anthony D. Long, Andrew G. Clark, Brian P. Lazzaro

## Abstract

Many of the molecular mechanisms for antifungal immunity in *Drosophila melanogaster* have been defined, but relatively little is known about the genetic basis for variation in antifungal immunity in natural populations. Using two population genetic approaches, Quantitative Trait Locus (QTL) Mapping and Evolve and Resequence (E&R), we explored the genetics underlying *D. melanogaster* immune defense against infection with the fungus *Beauveria bassiana*. Immune defense was highly variable both in the recombinant inbred lines from the Drosophila Synthetic Population Resource used for our QTL Mapping and in the synthetic outbred populations used in our E&R study. Survivorship of infection improved dramatically over just 10 generations in the E&R study, and continued to increase for an additional 9 generations, revealing a trade-off with uninfected longevity. Populations selected for increased defense against *B. bassiana* evolved cross resistance to a second, distinct *B. bassiana* strain but not to bacterial pathogens. The QTL mapping study revealed that sexual dimorphism in defense depends on host genotype, and the E&R study indicated that dimorphism also depends on the specific pathogen to which the host is exposed. Both the QTL Mapping and E&R experiments generated lists of potentially causal candidate genes, although these lists were non-overlapping.

## Introduction

Studies of insect immune defense have focused predominantly on immune mechanisms against bacteria and viruses, while defense against entomopathogenic fungi remains poorly understood. Dissecting the genetic basis for variation in insect susceptibility to fungal entomopathogens has potential to guide biological control efforts. *Beauveria bassiana* is an entomopathogenic fungus that has been used to control crop pests that threaten food security for a growing international human population (Ugine et al. 2005, Li et al. 2010). Fungal biocontrol also has major prospective public health impact through suppression of disease vector insects. For example, *B. bassiana* can be deployed for management of bed bugs (Barbarin et al. 2012) and has potential to limit mosquitos in the genera *Aedes* and *Anopheles* (Garcia-Munguia et al. 2011, Valero-Jiménez et al. 2017). Susceptibility to *B. bassiana* is genetically variable within populations of insects (Tinsley et al. 2006). This variability can provide the substrate for natural selection to increase resistance, which may thwart control initiatives. Given the broad host range of *B. bassiana*, this pathogen is unlikely to be in any strict coevolutionary arms races with its varied hosts. Thus we can effectively use the fruit fly, *Drosophila melanogaster*, which shares many homologous immune defense genes and pathways with other insects as a model for defense against this generalist fungus.

Studies with *D. melanogaster*, have already identified several genes and pathways involved in insect immune defense against fungal pathogens (Buchon et al. 2014), but much of the inter-individual variation in immune defense against fungal infection still remains unexplained. Defense against the generalist *B. bassiana* likely involves many genes, and their interactions with each other and with the environment (Lu et al. 2016). Understanding the genetic basis for immune defense is complicated by pathogen exposure, diet, and other environmental factors. Even when these factors are controlled in the laboratory, identifying causative genetic variants for immune defense is challenging given that this trait is highly multigenic and can evolve quickly (Sackton et al. 2007) under positive selection (Schlenke and Begun 2003) or purifying selection (Han et al. 2013; Jiggins and Hurst 2003).

Here we used two approaches for identifying causative loci underlying variation in *D. melanogaster* defense against *B. bassiana*: QTL mapping, and evolve and resequence (E&R). For our QTL mapping study, we used a large multiparental advanced generation reference panel called the Drosophila Synthetic Population Resource (DSPR) (http://FlyRILs.com). The DSPR panel allows the estimation of effects of alleles that are at low frequency in natural populations, as long as these variants are present in the initial founder lines (Macdonald and Long 2007). Moreover, the DSPR has high power and high mapping resolution (King et al. 2012a). Thus using the DSPR, we potentially can identify subsets of rare alleles with large effects that influence immune defense. The DSPR have already been used to examine the genetic bases for various traits (reviewed by Long et al. 2014), such as caffeine resistance (Najarro et al. 2015), boric acid resistance (Najarro et al. 2017), and methotrexate toxicity (Kislukhin et al. 2013). We tested 296 recombinant inbred lines from the DSPR for survival after infection with *B. bassiana*, and identified several genomic positions correlated with variation in this trait. Males and females survived infection differently, with the direction and magnitude of sexual dimorphism being dependent on fly genotype.

In natural populations, alleles with large effects on *D. melanogaster* immune defense, of the type that can be identified via QTL mapping, may be selected against due to pleiotropy or trade-offs with other fitness characters. Thus, it is unclear whether there is substantial rapidly-selectable variation in defense against *B. bassiana* in natural *D. melanogaster* populations. Therefore, we used laboratory selection to test whether populations of *D. melanogaster* can evolve rapidly to become more resistant to *B. bassiana* and identified the alleles that changed in frequency over the course of this evolution. To begin the selection experiment, we used a large outbred laboratory-adapted population that was derived from mixing the Global Diversity Lines (Greenberg et al. 2010). Following ten and nineteen generations into selection for defense against *B. bassiana*, we tested control and selected populations for their susceptibility to infection with the selective agent, *B. bassiana* ARSEF 12460. To test for the evolution of cross-resistance against other pathogens, we also tested for changes in survival of infection with a second fungal pathogen (*B. bassiana* GHA), a Gram-negative bacterium (*Providencia rettgeri*), and a Gram-positive bacterium (*Enterococcus faecalis*). Selected populations rapidly evolved increased survival of infection with ARSEF 12460 compared to control populations. However, this improvement in immune defense came at a cost to uninfected longevity, demonstrating the existence of an evolutionary trade-off. Comparing Single Nucleotide Polymorphism (SNP) frequencies in the selected and control populations identified significantly differentiated sites, enriched in genes involved in multiple functional categories. Cross-resistance tests showed that selection for defense against *B. bassiana* ARSEF 12460 resulted in improved defense against *B. bassiana* GHA but did not affect resistance against bacteria.

## Materials and Methods

### 1. Quantitative Trait Locus Mapping

#### The Drosophila Synthetic Population Resource (DSPR)

We phenotyped 296 RILs from the A1 population of the DSPR (King et al. 2012a,b). For at least 8 generations prior to being phenotyped for immune defense, the RILs were moved to Cornell Biotech Glucose medium (per liter of deionized water: 82 g glucose, 82b Brewer’s yeast, 10g agar, 10 mL acid mix composed of 4.15% phosphoric acid by volume).

#### Preparation of RILs for phenotyping

Prior to each assay, 100 male and 100 female flies per RIL were anesthetized with carbon dioxide and placed in a bottle with a small Petri dish of medium supplemented with yeast paste (∼1 teaspoon yeast mixed in 1ml DI water). After 24 hours eggs were separated into vials at densities of 60-80 eggs/vial. Starting on day 13 from egg, flies were transferred to fresh vials daily until inoculation. Each RIL was tested in three biological replicates with the exception of 13 RILs that were tested only twice. To ensure that the timing of the test did not confound our results, we randomized the timing of the tests: a random “*group”* of the RILs underwent egg collection on the same day. The three replicates tested for each RIL were tested in different *groups* (with the exception of 36 RILS for which some replicates were tested in the same group). A random “*set”* from each *group* was inoculated when the RILs were 16 days old from egg and a second *set* was inoculated the next day. Thus, RILs in the same “*group*” were reared together, and RILs in the same *set* (nested within group) were sprayed together. The division into groups and sets was necessary for handling because 878 inoculation assays were performed in total.

#### Inoculation of flies with *B. bassiana*

Flies were inoculated with *B. bassiana* ARSEF 12460 Shahrestani & Vandenberg (Shahrestani et al. 2018). Flies were briefly anesthetized with carbon dioxide (CO_2_) and measured in a microcentrifuge tube to 0.5 mL, which corresponds to approximately 50 flies/sex. These flies were then spread out on a small Petri plate lid placed on ice to sustain anesthetization. Flies were sprayed with 5 mL of a fungal suspension (0.034 g spores/25 mL of 0.03% Silwet) using a Spray Tower (Vandenberg et al. 1996), which introduced approximately 100 spores/mm^2^ of *B. bassiana* to the fly cuticle. Inoculated flies were placed into cages and kept at 25° C and 100% humidity for 24 hours to allow the fungus to germinate. Afterward, the cages were maintained at 25° C at 60-70% humidity with the usual 12/12 hour-light/dark cycle. Mortality was counted daily and recorded for ten days, distinguishing the number of males and females that were dead or lost due to handling. After ten days, the surviving flies were terminated and counted to determine the exact number of flies that were in each cage.

#### Fungal Viability Check and Spore Count

To confirm spore viability a 2 mL suspension of a 1:1000 dilution of 0.34 g of lab grown *B. bassiana* in 25 mL of 0.03% Silwet was sprayed through the spray tower onto a 60 x 15 mm water agar Petri dish which was incubated at 25°C for 24 hours. Following incubation one hundred spores within a central swath were inspected under a light microscope for presence or absence of a growing germ tube. Spores with a germ tube greater than or equal to the length of the spore were tabulated as living while others were considered non-viable. The ratio of living vs non-living spores was used to determine % viability. Poor viability (less that 90%) required production of a new batch from the isolate from the ARSEF culture collection. Dried spores were stored at minus 20°C until checked for viability or dosing of insects. To estimate deposition of spores on the arena a plastic microscope cover slip (22 x 22 mm) was placed adjacent to each group of insects sprayed with a dose. After the cover slips were dried they were placed in a 50 mL centrifuge tube with approximately 15 small glass beads and 5 mL of 0.03% Silwet solution. A vortex shaker was used to dislodge spores from the coverslip into suspension (Ugine et al. 2005). Using a pipette, a drop of spore suspension was placed onto two hemocytometers. These four counts were used to estimate number of spores per ml deriving the number of spores/mm_2_ deposited on the fly spray arena.

#### Data Analysis

All data analysis was carried out using the R statistical programming language (R Core Team 2017). The focal phenotype for each RIL is the proportion surviving the fungal infection 10 days post-inoculation. The distributions of phenotypes were slightly skewed (Figure S1). Thus, to improve normality, we used an arcsine square root transformation, commonly used to transform proportion data (Figure S1). Using the lmer function in the lme4 package (Bates et al. 2015), the effects of group and set were tested by comparing mixed effects linear models that included group and set to models without each effect using the anova function. Set had no effect on either sex (males: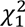 = 1.01, P = 0.31; females: 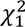 = 1.93, P = 0.16). Group was marginally significant in males but not significant in females (males: 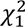 = 4.13, P = 0.04; females: 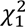 = 2.16, P = 0.14). But given the weak evidence for a group effect and the fact that for the vast majority of RILs, replicates were distributed among three groups, we did not include group or set in any future analyses. Using the same approach, line and sex did significantly affect the result and were considered for QTL mapping and estimation of heritability. We used the lme function in the nlme package (Pinheiro et al. 2020) to fit a mixed model ANOVA and used VarCorr to estimate variance components to estimate broad-sense heritability of the proportion surviving in males and females.

#### QTL mapping

The methodology for mapping QTL in the DSPR, and in multiparent populations more generally, has been described extensively previously (King et al. 2012a,b; Broman and Sen 2009). Briefly, a hidden Markov model assigns the underlying founder ancestry to each segment of each RIL with an associated probability (King et al. 2012b). At each of ∼10,000 positions across the genome, we regressed the mean line phenotype on the eight founder probabilities, analyzing males and females separately. We also mapped the difference between male and female survivorship to examine dimorphism in susceptibility. We used a permutation test (Churchill and Doerge 1994) to identify the genome wide significance threshold associated with a 5% family-wise error rate and to estimate the false discovery rate (FDR) (Benjamini et al. 1995). For each permutation, we calculated the average number of false QTL across all phenotypes at different significant thresholds. We first identified all peak positions for a given genome scan, then we removed any peaks that were within 2 centiMorgans of a higher peak. We then calculated the threshold that corresponded to the FDR. Here the FDR is the expected number of false positives/the expected number of total positives at a threshold of 5% and 50%. We used FlyBase (www.flybase.org) to convert the 5.x coordinates given by the QTL analysis to the updated 6.x coordinates. Then we used FlyBase (www.flybase.org) to identify the genes located in these regions of interest.

### 2. Experimental Evolution and Resequencing

#### *Drosophila melanogaster* populations

The base population was obtained from co-author AGC and was originally created by combining ninety-six isofemale lines from five geographic areas (Greenberg et al. 2010). These were round-robin crossed (first per population, then across populations), allowing for populations to become genetically diverse over time. The lines were from Beijing (Begun and Aquadro 1995), Netherlands (Bochdanovits and Jong 2003), Ithaca NY, Tasmania, and Zimbabwe (Begun and Aquadro 1993). The outbred population was maintained on discrete 14-day generations in a 12:12 light/dark incubator, with developmental phase in bottles and egg-laying phase in cages.

#### *Beauveria bassiana* inoculations and verification of dose and viability

For immune defense selection, we used *B. bassiana* ARSEF 12460 (Shahrestani et al. 2018). Specifically, 0.34 g of *B. bassiana* spores were suspended in 25 ml of 0.03% Silwet in DI water. This suspension was then diluted 1:10, 1:100, and 1:1000 for the dose response assay. For each inoculation, 7.5 ml of suspension was sprayed onto flies and the viability and dose of *B. bassiana* were estimated using the protocols described above. The selection dose (undiluted) was ∼10^4^ spores per mm^2^, and the fungus was fully viable in all sprays. In each infection assay, some of the flies that died were plated on SDAY media and monitored for fungal sporulation on their cuticle, thus showing that dead flies had been infected. Control groups underwent the same treatment as the inoculated groups, but were sprayed with just 0.03% Silwet suspension (no fungus). Cadavers from the control groups did not show sporulation on their cuticle.

#### Selection Protocol

After growing the ancestral population size over six generations, we divided it into eight groups and randomly assigned these groups to four control (C_1-4_) and four selected populations (S_1-4_). For each of the eight populations, 10,000 eggs were collected per generation. In the S_1-4_ populations, these 10,000 flies (estimated by volume: 2,000 flies was approximately 5ml of volume) were inoculated with *B. bassiana* and then divided into five cages at densities of 2,000 flies per cage. For each of the four C populations, 10,000 eggs were collected and after eclosion, 2,000 adults were randomly chosen and sprayed with 0.03% Silwet and placed into a single cage. Dead flies were removed from the cages daily, preventing secondary infection from cadavers. Flies were fed daily with Petri plates filled with medium. When 80% of the flies in the S populations died, surviving flies across all five cages of each replicate population were combined to form a single cage with approximately 2,000 flies and given yeast supplement in addition to the diet to promote oviposition. Over the following 1-3 days, 100 vials of eggs at densities of ∼60-80 eggs/vial were collected for each population to yield over 10,000 eggs. This protocol was repeated for 19 generations. Each C_i_ population was kept on the same timing as its corresponding S_i_ population.

#### Infection resistance assays

At generations 10 and 19, we compared the C_1-4_ and S_1-4_ populations for divergence in resistance to four pathogens: *B. bassiana* ARSEF 12460, *B. bassiana* GHA, *Providencia rettgeri* (Gram-negative bacterium), and *Enterococcus faecalis* (Gram-positive bacterium). For the two fungal pathogens and for their matched uninfected control, the sample sizes were ∼100 flies per sex per population, tested in two replicate cages. Flies were handled in the same manner as in the selection protocol, except they were kept in smaller cages to control for density effects, and dead flies were removed and sexed daily until all flies died. For the two bacterial pathogens the sample sizes were ∼100 flies per sex per population, and ∼40 flies per sex per population for the wounding control groups. These flies were anesthetized in groups of 15 or fewer on CO_2_ and pricked in the thorax with a needle dipped in dilute bacterial culture, or with a sterile needle as a wounding control, and then maintained in groups of ten in vials (Khalil et al. 2015). Bacterial cultures were grown overnight in Luria broth at 37°C from a single bacterial colony, and then diluted with sterile LB to O.D._600_ = 1.0 (for *P. rettgeri*) or O.D._600_ = 0.5 (for *E. faecalis*).

#### Statistical analyses for phenotypes

The Kaplan-Meier estimate of the survival function (Kaplan and Meier 1958) and tests of significance using Cox Proportional Hazard model (Cox and Lewis 1972) were performed using the package Survival in R. Survival after infection with *B. bassiana* ARSEF 12460 was analyzed with the following model:

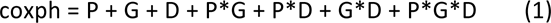

where * shows interaction, G represents generation (10 and 19), P represents populations (Control and Selected), and D represents the infection status (uninfected and infected with 10^4^ spores/mm^2^). The variables of population replicate, treatment replicate, and sex, did not significantly affect hazard ratios in preliminary analyses and were omitted from the final model. To study the effect of dose when infected with *B. bassiana* ARSEF 12460, we looked at the LT50, the median lethal time in days, and investigated the effect of population (S vs. C), sex, generation (10 vs. 19), and dose on LT50 with ANOVA.

Survival after infection with *B. bassiana* GHA was analyzed with the following model:

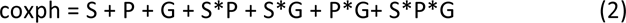

where the notation is the same as for model (1), with the addition of S, which represents sex (female and male). Keeping the three-way interaction term, which was borderline significant, improved the validity of the proportional hazard, and hence the fit of the model. Since the interactions of sex with other factors were significant, though the marginal effect of sex was nonsignificant, we kept this term in the model to follow a hierarchical modeling approach. Unlike with ARSEF 12460 and GHA, no difference between S and C populations was observed for resistance to the two bacterial infections (*E. faecalis* and *P. rettgeri*). The hazard ratios in all sub-populations were 1. The p-values of log-rank tests to compare survival functions in each sub-group were very high, confirming that Hazard Ratio (HR)=1 across the board. Therefore, no further analysis was conducted on these data.

#### DNA Extraction and Sequencing

After zero and 19 generations of selection, samples of adult female flies were frozen in a −80 freezer. Nine genomic DNA libraries were made from these samples, each with 100 frozen females (pooled): one from each of the C_1-4_ and S_1-4_ populations and one from the founding population. These were sequenced on Illumina Hi-Seq platforms within the Cornell sequencing core facility.

#### Mapping of Reads

Reads were mapped against the *D. melanogaster* reference genome (version 6.14) using BWA (version 0.7.8) (Li and Durbin 2009) using bwa mem with default settings. We filtered and sorted the resulting SAM files for reads mapped in proper pairs with a minimum mapping quality of 20 and converted them to the BAM using the view and sort commands in SAMtools (Li et al. 2009). The rmdup command in SAMtools was then used to remove potential PCR duplicates. Average coverage was above 30X or greater for all populations except C3, which was 25X (Table S1). Next, bam files for all 9 populations were combined into a single mpileup file once again using SAMtools. The mpileup file was in turn converted to a “synchronized” file using the PoPoolation2 software package (Kofler et al. 2001). This file contains allele counts for all bases in the reference genome for each population in a succinct tab delimited file. Lastly, RepeatMasker 4.0.3 (http://www.repeatmasker.org) was used to create a gff file containing simple sequence repeats found in the *D. melanogaster* genome version 6.14. PoPoolation2 was then used to make these regions within the sync file.

#### Patterns of SNP Variation

A SNP table was created using the sync file described above. We only considered sites where coverage was between 15X and 200X, and for a site to be considered polymorphic we required a minimum minor allele frequency of 2% across all 9 populations. All sites failing to meet these criteria were discarded. Prior to performing any of the analyses described below, we sought to identify cases where a given nucleotide was fixed across all of the S populations but not the ancestral P populations and C populations, which may be expected for strong selection on standing variants. To assess broad patterns of SNP variation in P, C, and S populations, heterozygosity was calculated and plotted over 100 kb non-overlapping windows directly from the major and minor counts in our SNP table. A t-test was also performed to compare mean heterozygosity between the C and S populations. To assess how closely replicate populations resembled one another within the C and S groups, *F_ST_* estimates were also obtained using the formula: *F*_ST_= (*H*_T_-*H*_S_)/*H*_T_ where *H_T_* is heterozygosity based on total population allele frequencies, and *H_S_* is the average subpopulation heterozygosity in each of the replicate populations (Hedrick 2011). *F*_ST_ estimates were made at every polymorphic site in the data set for a given set of replicate populations.

#### SNP Differentiation

We used two different methods to characterize SNP differentiation between the C and S populations. First, we used the Cochran-Mantel-Haenzsel (CMH) test as implemented in the PoPoolation2 software package to compare SNP frequencies at every polymorphic site in our SNP table between the two groups of populations. CMH tests between our two groups of populations were performed at each of these polymorphic sites. To establish a significance threshold for these tests, we first performed simulations to generate a distribution of P-values associated with a null expectation of genetic drift rather than selection (See “Simulations” section below). Briefly, we generated sets of 8 populations based on allele frequencies in the ancestral P populations, then simulated 19 generations of genetic drift. Within each set of 8 populations, half were randomly assigned as “control” and the remainder were randomly assigned as “selected.” CMH tests were then performed at each polymorphic site between the two groups. This was done 100 times and all the resulting p-values were recorded. The quantile function in R was then applied to these p-values to define a significance threshold that corresponds to a genome-wide false-positive rate, per site, of 5%. This ultimately resulted in a significance threshold of approximately 3.94 × 10^-18^.

Along with the CMH test, we also compared SNP frequencies between the C and S populations using the quasi-binomial GLM approach suggested by Wiberg et al. (2017). Based on their findings, this approach is reported to have lower false positive and higher true positive rates than the CMH tests. However, it should be noted that our simulation-based approach to correcting for multiple comparisons when using the CMH test resulted in a more stringent significance threshold than what was featured in their work. The quasi-binomial GLM test was implemented using scripts provided by Wiberg et al. (2017). As suggested by the authors, allele counts were scaled to the effective sample size (*n_eff_*) as described in Kolaczkowski et al. (2011) and Feder et al. (2012). As counts of zero can lead to problems when implementing this approach (see Wiberg et al. 2017), a count of 1 was added to each allele whenever a zero was encountered. In terms of correcting for multiple comparisons, one of the reported benefits of quasi-binomial GLMs is that they produce the expected uniform distribution of p-values under the null hypothesis which allows for standard methods of correcting for multiple comparisons (Wiberg et al. 2017). To that end, instead of the simulation approach used for our CMH tests, we opted to use a Bonferroni correction, and the q-value approach (Storey et al. 2017, Storey and Tibshirani 2003). We chose these two methods as Wiberg et al. (2017) found them to be the most and least conservative approaches, respectively.

#### Gene Ontology (GO) Analysis

The Gowinda software package (Kofler and Schlotterer 2012) was used to identify enriched GO terms based on our candidate sites. Our list of candidate sites consisted of the 45 significantly differentiated SNPs identified between the C and S populations based on our CMH comparison. The background list contained our complete SNP list based on our previously described SNP calling parameters. A gene annotation file for the *D. melanogaster* reference genome (6.14) was obtained from FlyBase, and a gene set file for relevant GO terms was obtained from FuncAssociate3 (Berriz et al. 2009). With these inputs, Gowinda was set to run for 10^6^ simulations with the gene-definition and mode parameters set to “gene”. This analysis identified 132 GO terms with p-values < 0.05. This list was then filtered so all GO categories containing less than 2 reference genes were discarded, and the resulting list was run through GO-Module to correct for hierarchical clustering (Yang et al. 2011).

#### Simulations

To perform our genetic drift simulations, we used MimicrEE (https://sourceforge.net/projects/mimicree/), a forward simulation specifically designed to mimic experimental evolution. MimicrEE simulates populations of diploid individuals where genomes are provided as haplotypes with two haplotypes constituting a diploid genome. There are no changes in the demography once the initial population file is submitted and a list of selected loci may be provided. The simulated populations have non-overlapping generations and all individuals are hermaphrodites (though selfing is excluded). At each generation, matings are performed, where mating success (number of offspring) scales linearly with fitness, until the total number of offspring in the population equals the targeted population size (fecundity selection). Each parent contributes a single gamete to the offspring. As we were only interested in simulating genetic drift, we did not specify any fitness differences between different genotypes. Crossing-over events are introduced according to a user-specified recombination rate. The recombination rates were specified for 100 kb windows and were obtained from the *D. melanogaster* recombination rate calculator v2.2 (Fiston-Lavier et al. 2010). As recombination does not occur in male *D. melanogaster*, the empirically estimated female recombination rate was divided by two for the simulations.

The starting populations used in our simulations were generated based on SNP frequencies in the ancestral P population across the 268,272 polymorphic sites along chromosome 3R. Each starting population consisted of 600 individuals. To create each individual’s genotype, two random numbers in the range [0.0, 1.0) were generated at each polymorphic site. These numbers were then compared to the ancestral data’s major allele frequency at the position. If the random number was less than the major allele frequency, the major allele was added. Otherwise, the minor allele was added. In this manner, we generated 100 sets of populations each consisting of 8 populations derived from the SNP frequencies in the P populations. All sets of populations were then subjected to 19 generations of drift. Within each set, the populations were then randomly split into two groups of 4 and the CMH tests were performed at each polymorphic site between the two groups. All p-values were recorded and, the quantile function in R was used to define a significance threshold that corresponds to a genome-wide false-positive rate, per site, of 5%.

We estimated effective population size (*N_e_*) in our experimental populations using the Nest package in R, which was specifically designed to estimate *N_e_* from temporal allele frequency changes in experimental evolution (Jonas et al. 2016). This analysis suggested *N_e_* to be ∼600, so we simulated populations of 600 individuals. As genotype inputs, we used SNP frequencies in P populations for the first time point and frequencies in the C populations for the second. We used the “P.planI” method for estimating *Ne*, which was specifically to account for the two stage sampling process associated with pool-seq data (individuals being sampled from a population, and reads being sampled from DNA pool). *Ne* estimates varied depending on which of the C populations was used, but the average was 592 or ∼600 (N_e_ was 466 for C_1_, 616 for C_2_, 590 for C_3_, and 696 for C_4_).

#### Data sharing plan

All relevant data and script files were uploaded to Figshare along with descriptions. These files include raw data for the RIL phenotypes, SNP tables, CMH and GLM results, and major scripts.

## Results

### 1. Quantitative Trait Locus Mapping

We measured survival ten days after inoculation with *B. bassiana* in 297 RILs from the DSPR. Survival ranged from 0 to 92.25% (Figure 1). For males, mean survival was 58.97% and median survival was 63.77%, and for females the mean and median survivals were 52.45% and 53.93% respectively. The replicates tests for each RIL were similar to each other (see standard errors of the means in Figure 1); the RIL with the large differences among replicates was RIL11039, which may have been more affected by slight environmental differences among replicates. Heritability of the median survival time was 0.88 for males and 0.85 for females.

**Figure 1.**
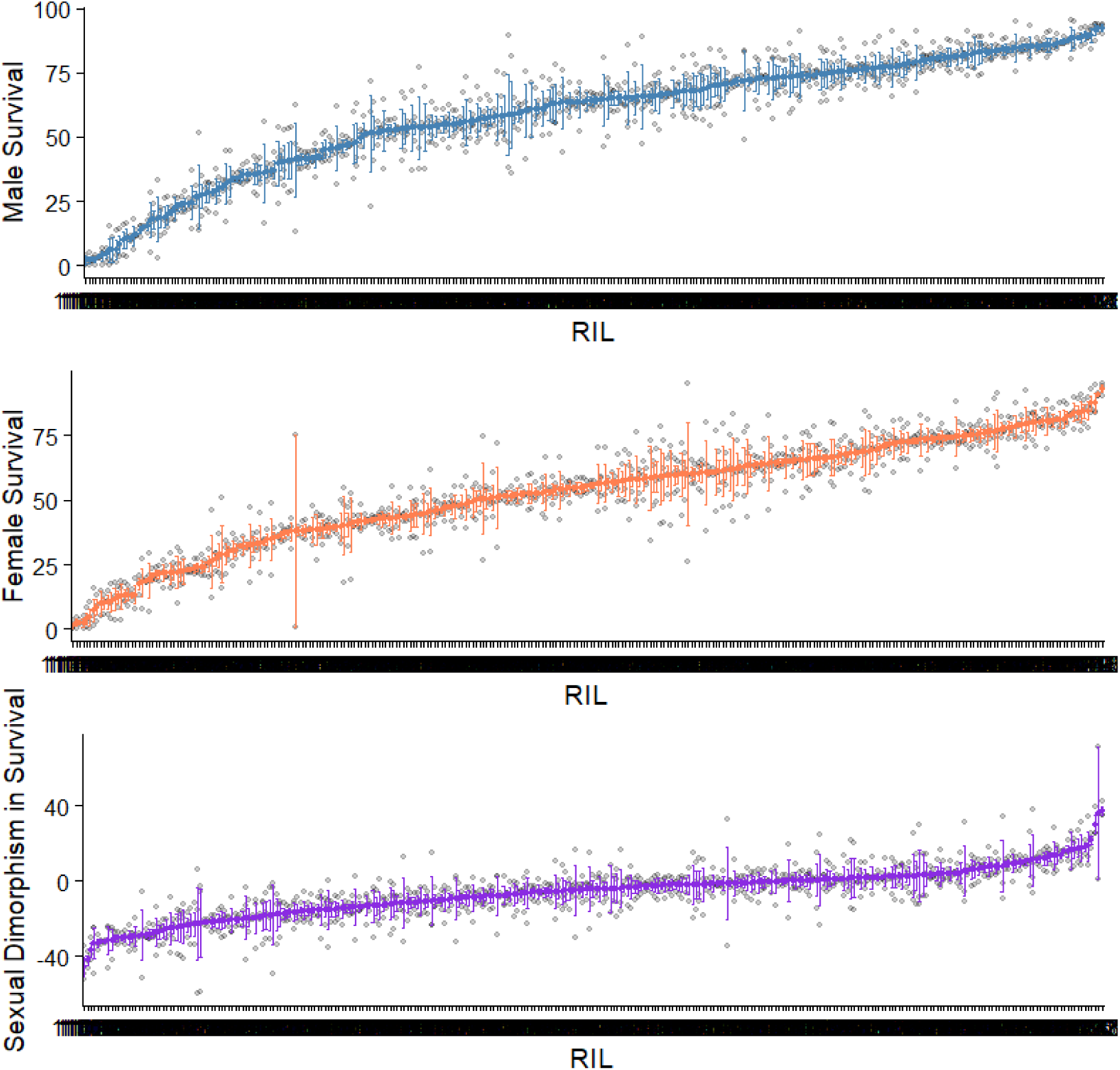
Survival of the DSPR RILs post-inoculation with *B. bassiana* ARSEF 12460. Survival 10-days post inoculation is shown for males (A) and females (B). Male survival subtracted from female survival gave a measure of sexual dimorphism in survival (C). In some RILs males survived better than females, in others females survived better than males, and in some there was no sexual dimorphism. Bars show standard error of the mean as each RIL was tested multiple times.

Most RILs had a sexually dimorphic response to infection, with males and females exhibiting different probabilities of surviving 10 days after infection (Figure 1). Moreover, the magnitude and even the direction of the sexual dimorphism varied across RILs (Figure 1). In 68.35% of the RILs, more females than males were dead 10 days after infection, and in the remaining RILs the direction of dimorphism was reversed (Figure 1). Within RILs, the direction of dimorphism remained highly consistent across replicates. Nevertheless, across RILs, immune defense was positively correlated (*r*^2^ = 0.59; p < 2.2×10^-16^) between males and females indicating sex-independent genetic differences among the lines (Figure S2).

The family-wise significance threshold for QTL was a LOD score of 8.07, which was similar to the 5% FDR threshold (8.1), and using either yielded the same set of QTL. We identified one significant QTL using these thresholds from the male data (Figure 2). We additionally considered a more liberal FDR of 50%. At this lower threshold, we identified three QTL peaks (one peak for males, one for females, and one for the dimorphism) (Figure 2). The precise coordinates of these peaks are shown in Table S2.

**Figure 2.**
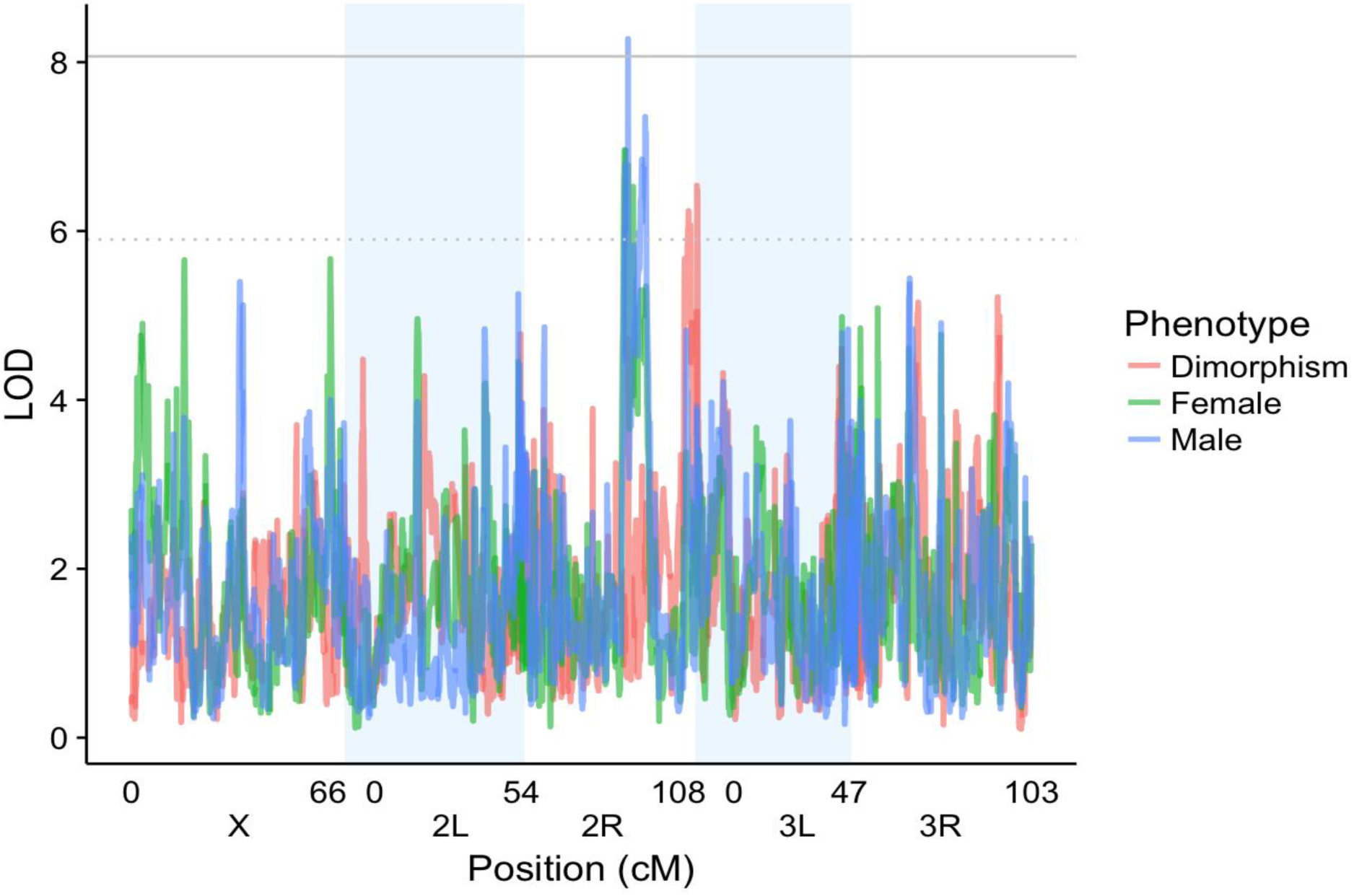
Positions of loci contributing to variation in immune defense in the QTL mapping study. The LOD score is plotted for each genomic position across chromosomes X, 2, and 3. The solid grey line is the 5% FDR and FWER (they are very close to each other). The dotted grey line is the 50% FDR. The QTL Mapping was done separately for males (blue), females (green), and the sexual dimorphism (red).

There were 456 genes within the peaks identified in our QTL mapping, 27 of which were under the 5% FDR peak. We explored these genes in Flybase (www.flybase.org) for their functions (Table S3). The genes included protein-coding genes, some with functional information available on Flybase (Table S3).

### 2. Experimental Evolution and Resequencing

#### Defense against *B. bassiana* ARSEF 12460

The summary fit of the Cox Proportional Hazards model (see Methods model 1) is presented in the Table 1 (Schoenfeld residual plots, not shown, confirmed the validity of proportional hazard assumptions). Selection for defense against *B. bassiana* ARSEF 12460 resulted in improved survival of infection, with selected populations living nearly twice as long when infected compared to control populations (Figure 3, Table 2). But this improved survival of infection came at a cost to survival under pathogen-free conditions, such that populations that evolved better immune defense had reduced lifespan than controls when uninfected (Figure 3, Table 2). Notably, the outbred populations in this study did not show sexual dimorphism in survival without infection, nor when infected with *B. bassiana* ARSEF 12460. We compared the different groups’ survivals by estimating the hazard ratio, which is a statistical measure of relative chances of death at all ages. For example, in Table 2, the estimated hazard ratio of 0.46 for infected S vs. C at generation 10 means that the relative instantaneous probability of death for an infected selected fly was 46% of the instantaneous probability of death for an infected control fly, and the p-value for this comparison was p<0.00001, implying significant divergence between survival of infection in the S and C populations. In other words, after just 10 generations, selection had resulted in substantially better survival of infection in the S populations. This difference increased by generation 19, such that at the end of the experiment, the instantaneous probability of death for an infected fly in the S populations was only 27% of the probability of death for an infected fly in the C populations (p<0.00001) as estimated from a Cox Proportional Hazard Model. The hazard ratio (S vs. C) among infected flies in generation 10 was significantly different from that of infected flies in generation 19 (the p-value of the test H_0_: HR_Gen10_=HR_Gen19_ for infected groups was 0.00021), suggesting that between generations 10 and 19, the S populations continued to evolve improved survival of infection compared to the control populations.

**Figure 3.**
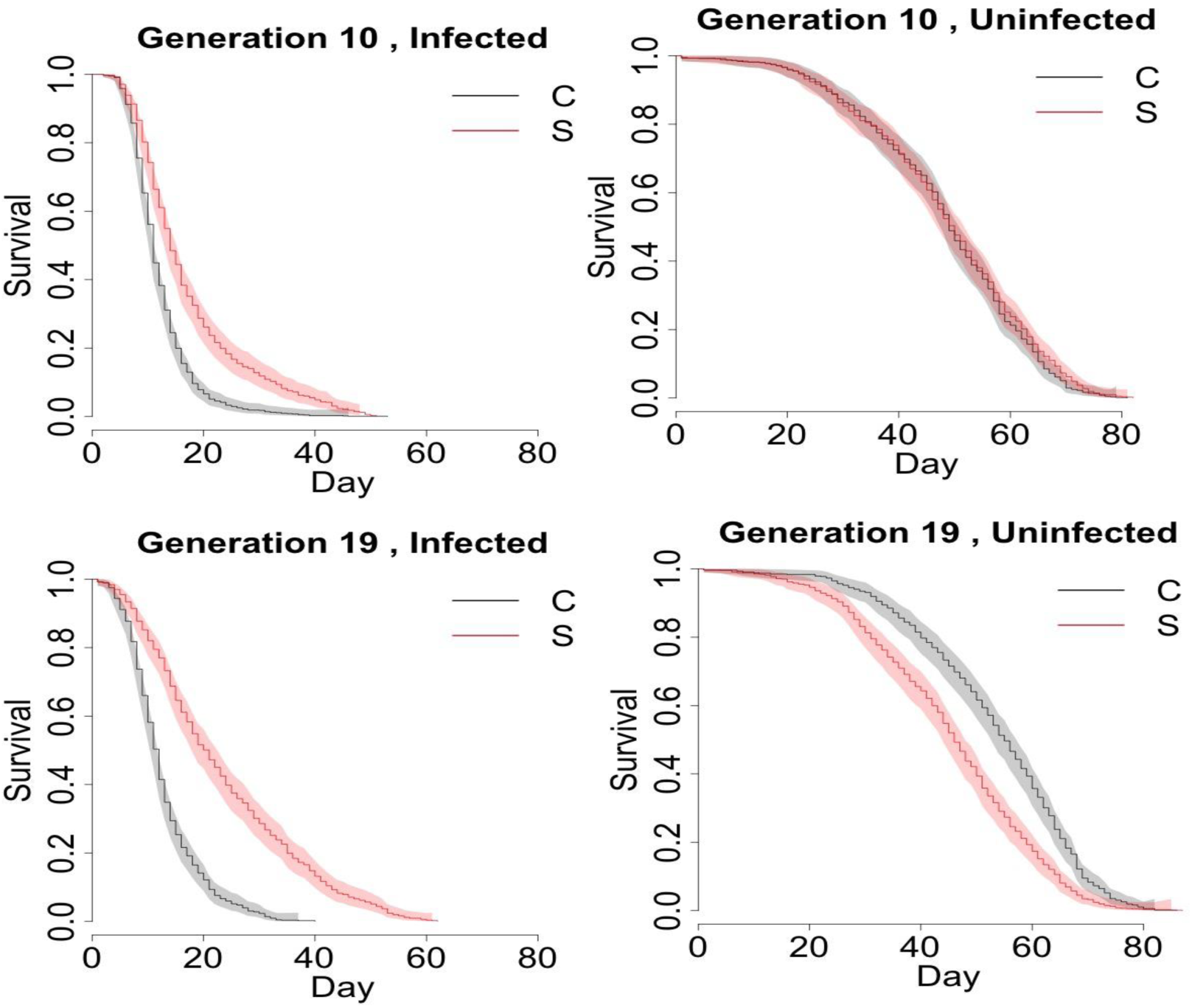
Survival of selected and control populations with and without infection with *B. bassiana* ARSEF 12460 at generations 10 and 19. The Kaplan-Meier estimate of the survival function is shown for S_1-4_ (red) and C_1-4_ (black) populations across different combinations of infection status (columns; infection with *B. bassiana* ARSEF 12460) and generation (rows). The shaded area represents 95% confidence intervals. After 10 generations of selection for defense against the fungal entomopathogen *B. bassiana* ARSEF 12460, the selected, S, populations had higher survival compared to the control, C, populations when infected with *B. bassiana* ARSEF 12460. After 19 generations of selection, the difference between infected S and C populations was even more pronounced. When the S and C populations were not infected, they did not differ in survival at generation 10, but at generation 19, the S populations survived worse compared to the C populations when uninfected, suggesting that their improved immune defense came at a trade-off with longevity in the absence of infection.

**Table 1.** Analysis of survival differences in selected and control populations with and without infection with *B. bassiana* ARSEF 12460 at generations 10 and 19. Survival was analyzed with model 1: coxph = P + G + D + P*G + P*D + G*D + P*G* where G represents generation (10 and 19), P represents populations (C and S), and D represents the infection status (uninfected and infected with 10^4^ spores/mm^2^). N= 7007.

|  | coef | exp(coef) | se(coef) | z | Pr(>\|z\|) |
| --- | --- | --- | --- | --- | --- |
| generation 19 | -0.341 | 0.711 | 0.049 | -6.982 | 2.91e-12 *** |
| population S | -0.086 | 0.918 | 0.049 | -1.760 | 0.078 |
| infection | 3.061 | 21.347 | 0.055 | 55.358 | < 2e-16 *** |
| generation 19: population S | 0.560 | 1.751 | 0.069 | 8.117 | 4.44e-16 *** |
| generation 19: infection | 0.240 | 1.272 | 0.067 | 3.582 | 0.000341 *** |
| population S: infection | -0.682 | 0.506 | 0.068 | -9.979 | < 2e-16 *** |
| generation 19: population S: infection | -1.113 | 0.328 | 0.097 | -11.516 | < 2e-16 *** |

**Table 2.** Hazard ratios from the Cox Proportional Hazard Model comparing selected (S) and control (C) populations across generation and infection status with *B. bassiana* ARSEF 12460. The reported p-values test the hypothesis H_0_: HR=1, or no difference in hazard between the S and C populations.

|  | Generation 10 |  | Generation 19 |  |
| --- | --- | --- | --- | --- |
|  | Infected | Uninfected | Infected | Uninfected |
| Hazard Ratio<br>(S vs. C) | 0.46 | 0.92 | 0.27 | 1.61 |
| p-value | < 0.00001 | 0.12 | < 0.00001 | < 0.00002 |

Interestingly, after 19 generations of selection, the instantaneous probability of death for an uninfected fly in the S populations was 161% of the instantaneous probability of death of an uninfected fly in C populations (Table 2; p<0.0002), meaning that populations that evolved improved immune defense paid the cost of reduced longevity in the absence of infection. This trade-off was not yet present at generation 10 (Table 2; p= 0.12). Thus, among uninfected flies, the hazard ratio (S vs. C) was lower at generation 10 than at generation 19 (the p-value of the test H_0_: HR_Gen10_=HR_Gen19_ for uninfected groups is 0.00083).

We investigated the effect of population (S vs. C), sex, generation (10 vs. 19), and dose on LT_50_ with an ANOVA model (Table S4; Figure S3). The LT_50_ increased with decreasing dose in every subgroup, but the rate of increase varied among subgroups (Figure S3). A simplified summary of the LT_50_ comparison between populations (S vs. C) for different doses is visualized in Figure 4, where we see a drop in LT_50_ for C populations from the uninfected (dose 0) group to lowest infection dose at 0.001, which is more drastic than the same drop in the S populations. The decrease in LT_50_ is more gradual in the S populations compared to C populations. This implies that not only did the S populations evolve increased resistance to infection, but they are also more robust to escalating dose of the pathogen. LT50 was slightly higher (by a factor of 10%) in generation 19 compared to generation 10. Males and females did not differ in LT_50_ in either the S or C populations, nor when all data were combined (Table S4; Figure S3).

**Figure 4.**
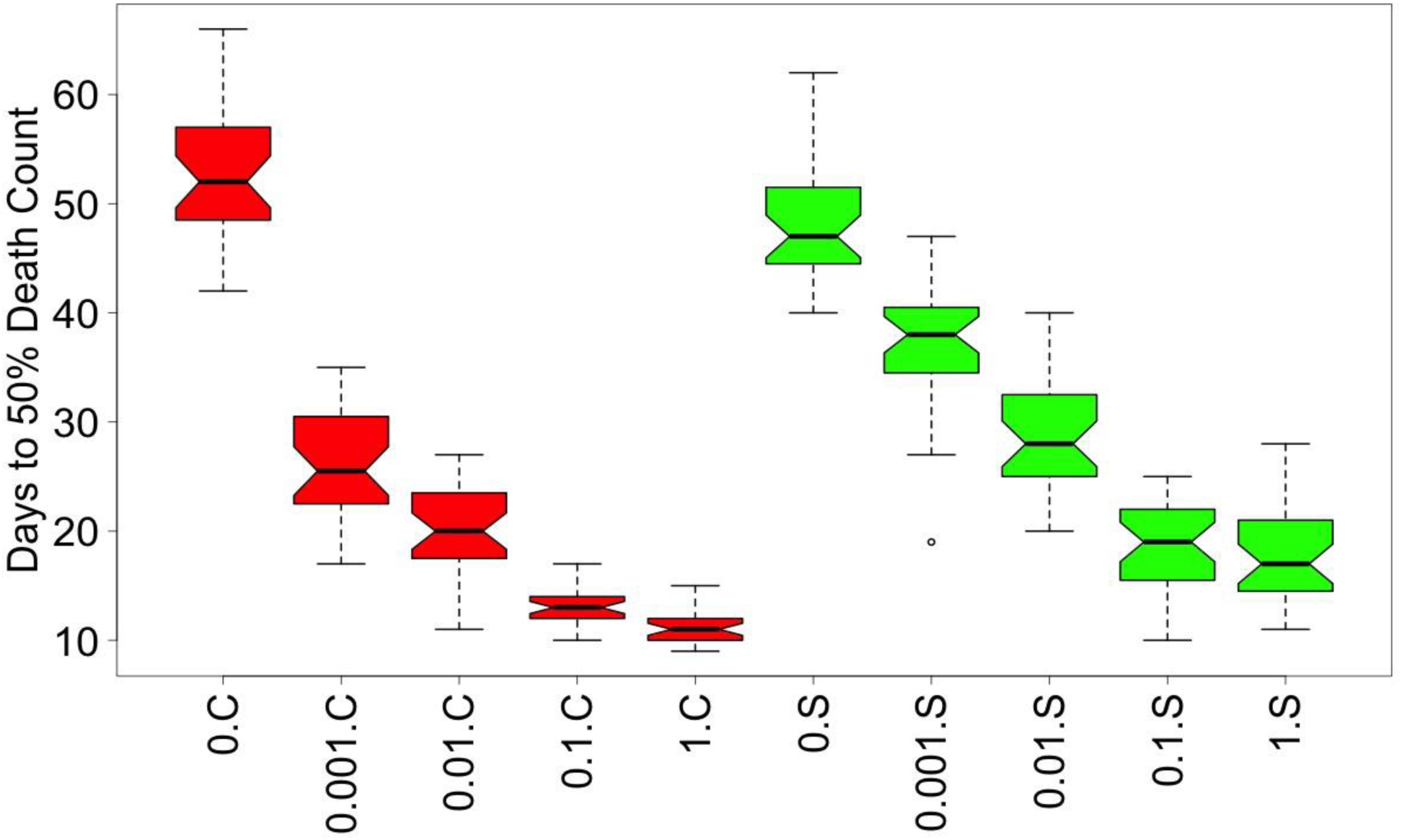
The median lethal time in days, or LT50, categorized by population type and infection dose. Data is combined across the replicate populations within each population type. Within each color, the 5 boxplots represent the 5 doses: 0, 0.001, 0.01, 0.1 and 1 (proportion relative to full dose of 10^4^ spores per mm^2^ of *B. bassiana* ARSEF 12460). LT50 was significantly affected by population (C vs. S, p <0.0001), generation (p=0.036) and dose (p<0.0001), but not by sex.

#### Defense against *B. bassiana* GHA

The summary fit for the Cox Proportional Hazards model (see Methods model 2) is presented in Table S5. All four replicate populations that were selected for resistance against *B. bassiana* ARSEF 12460 also became more resistant to a second fungal strain, GHA (Figure S4, Table S6). The magnitude of the S vs C difference in survival after infection with GHA depended on both sex and generation (Table S5). For males, at generation 10, the instantaneous probability of death for an S fly was 62% of the probability of death for a C fly (p= 0.0002) and by generation 19, the S males were surviving even better after infection, with an instantaneous probability of death of 44% compared to the C flies (p<0.0001). In females, the difference between C and S in post-infection survival was not significant at generation 10, with the probability of death of an S female being 82% of that for a C female (p=0.0243) but became significant by generation 19 with 45% probability of death of S vs. C (p<0.0001). This suggests that male flies evolved cross resistance to GHA faster than female flies. While overall survival after infection with GHA was higher in S flies than C flies at both generations 10 and 19 (Table S6), the magnitude of this difference was larger in generation 19 (Table S5)

Further supporting the sex differences in response to selection, we saw sexual dimorphism in the hazard ratio at generation 10 (H_0_: HR_male,Gen10_=HR_female,Ge10_, p-value = 0.0256) with males surviving better than females, but not at generation 19 (H_0_: HR_male,Gen19_=HR_female,Ge19_, p =0.7134). This difference in sexual dimorphism across generations can also be noted from the sex-by-population-by-generation interaction (p<0.0001; Table S5). Moreover, for both sexes, the hazard ratio (S to C) in generation 10 is statistically higher than that of generation 19 (H_0_: HR_male,Gen10_=HR_male,Ge19_, p = 0.0066, and H_0_ HR_female,Gen10_=HR_female,Ge19_, p = 0.0004). Selection for survival of ARSEF 12460 resulted in improved defense against GHA, but unlike with ARSEF 12460, defense against GHA was sexually dimorphic at generation 10.

#### Defense against bacterial pathogens

Unlike with fungal infection, no difference was observed between S and C populations for resistance to bacterial infections with *E. faecalis* and *P. rettgeri* after 10 or 19 generations of selection (Figure S5). There was also no difference between males and females (Figure S5). Flies infected with either bacterium were much more likely to die than uninfected (sterile pricked) flies at both generations 10 and 19 (log-rank tests p<0.0001). However, the likelihood of death was the same between S and C populations regardless of sex or generation (log-rank tests p>0.05).

#### SNP Variation

While we observe a number of chromosomal regions with notable depressions in heterozygosity in the P, C, and S populations, we do not see a dramatic loss in genetic variation in the C and S populations relative to the ancestral P population (Figure S6). Heterozygosity here is a quantification of population diversity, thus levels of variation are very similar in the C and S populations compared to the ancestral P population. This pattern is largely robust to changes in window size (Figures S7-S8). However, we do find that depressions in heterozygosity become more pronounced as window size is reduced. We find that mean heterozygosity at polymorphic sites in the ancestral P population is 0.24, and ranges from 0.23 to 0.24 in the C and S populations (Table S1). We do not find any significant difference in mean heterozygosity between the C and S groups (t-test p-value = 0.65). Among the C populations, we find that mean genome-wide *F_ST_* is 0.04. Mean genome-wide F_ST_ is also 0.04 among the S group. In both cases, this suggests a high degree of similarity between replicates of a given group. The fact that levels of *F_ST_* are the same in each group is also consistent with the duration of the experiment (i.e. there was not sufficient time for drift to produce high levels of divergence between replicates of a given group).

We examined sites that were fixed in all S populations at the end of the experiment but that remained polymorphic in the P and C populations. There were ∼4,000 such sites distributed across chromosomes X, 2 and 3 (Table S7). In about 98% of these instances, the frequency of the allele fixed in the S pops was ≥ 0.8 in the ancestral population. The lowest frequency we see in the ancestral population across all of these cases is 0.65. The SNP frequencies in the control populations at these ∼4000 sites follow the expectations from drift and have slight deviations from the ancestral frequencies. For comparison, we examined sites fixed in the C populations at the end of the experiment that were polymorphic in the S and P populations, which resulted in ∼1,000 sites (Table S8). These ∼1,000 sites had high (>0.8) frequencies in the ancestral populations.

#### SNP Differentiation

Our Cochran-Mantel-Haenzsel (CMH) tests comparing SNP frequencies in the C and S populations identified a total of 45 significantly differentiated SNPs across the major chromosome arms (Figure 5A). The majority of these sites were found on the X chromosome, while the remaining were split between 3R, 3L, and 2R. However, this pattern was not reflected in our quasi-binomial GLM results. Here no significantly differentiated sites were detected after the Bonferroni correction (Figure 5B) was applied, or when the less stringent q-value approach was used to correct for multiple comparisons (Figure 5C). Wiberg et al. (2017) report that the quasibinomial GLM approach has lower false positive and higher true positive rates than the CMH tests. However, given our simulation based approach for correcting for multiple comparisons when using the CMH test, we have a more stringent significance threshold than what was used in Wibert et al. (2017). Comparing Q-Q plots for the two approaches (Figure S9), the CMH test appears better suited for our data than the quasi-binomial GLM method (note the different y-axis ranges in the two Q-Q plots). SNPs and associated p-values from the three analyses are listed in Table S9.

**Figure 5.**
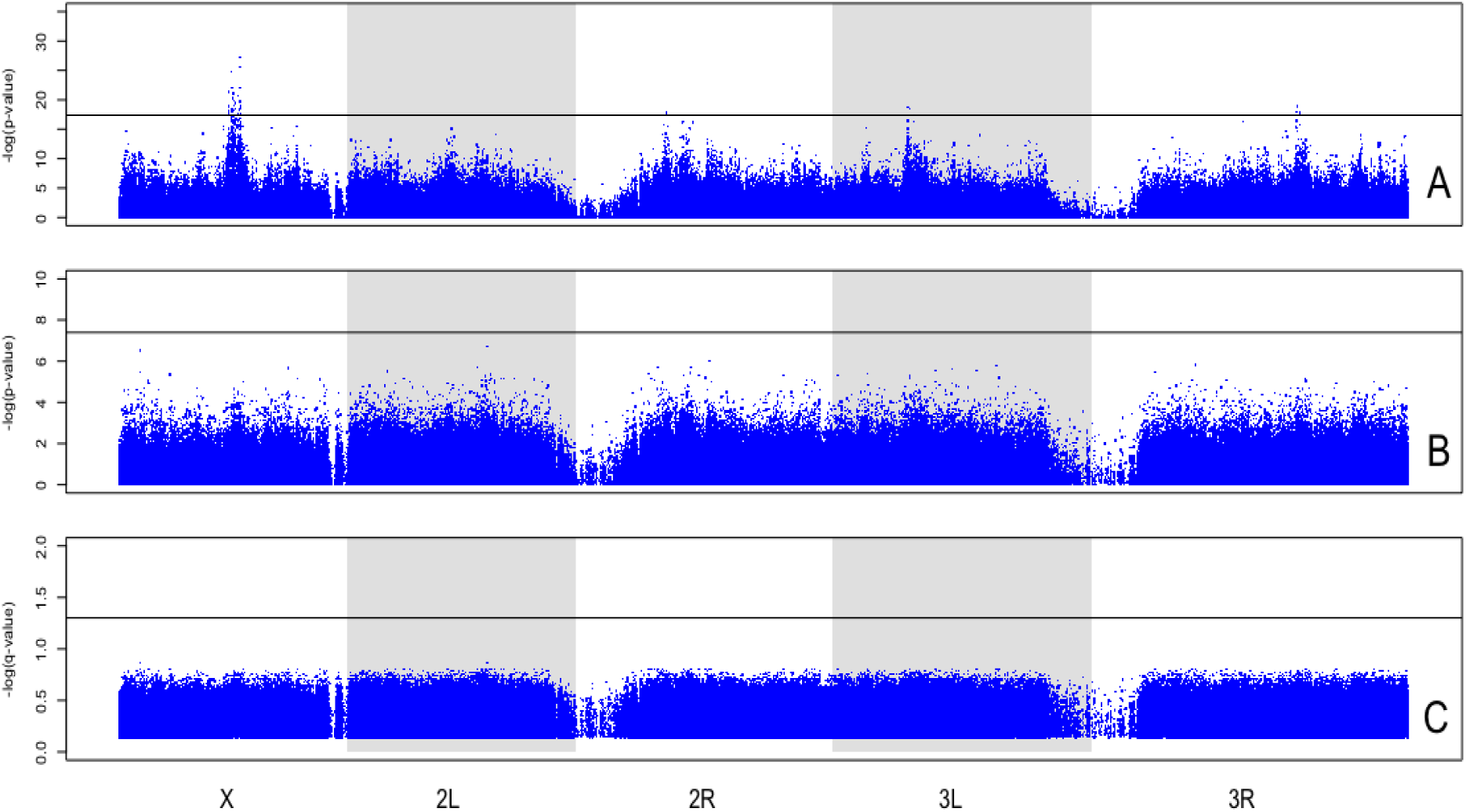
SNP frequency differences in the C and S populations. (A) Results from CMH tests plotted along all major chromosome arms as –log(p-values). Black line indicates Simulation derived significance threshold. (B) Results from quasibinomial GLM approach plotted as −log(p-values). Black line indicated Bonferroni corrected significance threshold. (C) Results from quasibinomial GLM converted to q-values, and plotted –log(q-values). A 0.05 false discovery rate threshold was used, as shown by the black line.

#### GO Terms

Running our list of candidate sites from the CMH tests through Gowinda, a tool that allows for analysis of gene set enrichment, identified 132 significantly enriched GO terms, which include nested and overlapping annotations (Kofler and Schlotterer 2012). Discarding terms with GO categories containing fewer than 2 genes and correcting for hierarchical clustering using GO-Module (Yang et al. 2011) reduced this list to 29 enriched terms (Table S10; See Table S11 for genes associated with each term).

## Discussion

Immune defense is a genetically complex and ecologically important trait with high levels of inter-individual variation. While much is already known about *D. melanogaster* immune defense against bacterial pathogens (reviewed in Buchon et al. 2014), defense against fungal pathogens has not been studied with the same depth. Yet insect defense against fungal infection has implications for biological control of crop pests and disease vectors. Here we applied two common approaches for mapping defense against a fungal pathogen to *D. melanogaster* genes, Quantitative Trait Locus (QTL) Mapping and Experimental Evolution and Resequencing (E&R) to characterize genetic variability in antifungal defense.

In our E&R approach, *D. melanogaster* populations that were exposed to *B. bassiana* infection quickly evolved improved defense against this fungus. There is some evidence that *B. bassiana* may be unique in terms of insect potential to evolve resistance to the pathogen under selection pressure (Dubovskiy et al. 2013). In our study, improvement in defense evolved within 10 generations and defense continued to improve until the termination of the experiment at nineteen generations. Rapid evolution of immune defense is possible because of standing genetic variation for host resistance. In wild populations, *D. melanogaster* defense against *B. bassiana* varies regionally (Tinsley et al. 2006; Paparazzo et al. 2015). For many traits, short-term adaptation appears to result primarily from existing genetic variants (Burke et al. 2010; Orozco-terWengel et al. 2012; Graves et al. 2017), and we would expect the same in our experiment. We maximized genetic diversity in our study by sampling flies from disparate geographic regions (Early and Clark 2017).

The evolution of immune defense came at a cost to uninfected longevity, suggesting an evolutionary trade-off with a genetic basis. Evolutionary trade-offs due to antagonistic pleiotropy may act to maintain genetic variation for life history traits and immune defense (Roff 2002; Schmid-Hempel 2003; McKean and Lazzaro 2011) and may constrain the evolution and maintenance of immune defense (Lazzaro and Little 2009). The large amount of variation for immune defense within natural populations suggests that this character commonly trades-off with other fitness components, and this has been experimentally observed in *Drosophila* (Kraaijeveld and Godfray 1997; Luong and Polak 2007, Vijendravarma et al. 2009, Ye et al. 2009). Such tradeoffs can result from genetic correlations between fitness traits and immune defense. In addition to antagonistic pleiotropy, genetic correlations among these traits may also result from linkage disequilibrium. There are high levels of inversions in some of the founder lines used in our study, but the extent to which the resulting linkage disequilibrium affected our observed phenotypes is unknown. Yet, some studies do not find any cost to laboratory evolved immune defense (Faria et al. 2015; Penley et al. 2018). For example, evolved immune defense through three selection regimes, oral and systemic infection with *P. entomophila* and systemic infection with Drosophila C virus, did not trade-off against reproductive output, development time, stress resistance, and other fitness characters (Faria et al. 2015), although Faria et al. (2015) did not examine potential trade-offs with uninfected longevity. It is possible that the costly trade-offs appear most prominently when there is a sudden shift to very high defense. For example, the selection pressure in the Faria et al (2015) study was much weaker than in our study, with 33% of their population surviving in the first generation and the percentage increasing in later generations due to adaptation. This idea is further supported by Duncan et al. (2011), who saw costs to the protozoan *Paramecium caudatum* that were selected for increased defense against the bacterial pathogen *Holospora undulata*. However, when selection was relaxed, the fitness was restored without completely losing the evolved resistance (Duncan et al. 2011). Therefore, there may be a threshold rate at which improved immune defense can evolve without noticeable fitness costs. This could explain why there were no fitness costs observed when *C. elegans* populations that were formerly selected for defense against *Serratia marcescens* maintained their increased resistance despite 16 generations of relaxed selection (Penley et al. 2018), or why no apparent fitness costs were seen when *D. melanogaster* were evolved for resistance to bacterial infection (Gupta et al. 2016).

The trade-off between immune defense and uninfected longevity is not axiomatic, because *D. melanogaster* populations that are experimentally evolved for resistance against other stressors, in particular starvation and desiccation, have increased longevity (Rose et al. 1992, Bubliy and Loeschcke 2005), and the increased longevity is sometimes maintained even after stress resistance reverts back to ancestral levels after a period of relaxed selection, even when the relaxed selection results from shifts in allele frequencies rather than any compensatory mutations (Phillips et al. 2018). Presumably, living longer requires effective stress resistance, and perhaps also a strong immune defense. Resistance and tolerance of infection both decline with age (reviewed in Garschall and Flatt 2018) and susceptibility to infection increases with age (Kubiak and Tinsley 2017), thus it may be expected that increases in longevity should be correlated with improvements in immune defense. Indeed, this has been observed, such that experimental evolution for delayed reproduction, which increases longevity, improves immune defense in *D. melanogaster* (Fabian et al. 2018).

Another potential reason, in addition to trade-offs with fitness characters, for populations maintaining variation in immune defense is that different genotypes are most resistant to different pathogens, which would lead to trade-offs within the immune system for defense against different pathogens. Evolution of immune defense against one pathogen may trade-off with defense against a second pathogen. For example, *D. melanogaster* selected for resistance against bacteria paid a cost in the presence of viruses (Martins et al. 2013). We tested our experimentally evolved populations for cross resistance against other pathogens, and found that evolution of immune defense against one strain of *B. bassiana* (ARSEF 12460), unsurprisingly, also led to improved defense against a second strain of *B. bassiana* (GHA). Fungi and Gram-positive bacteria are known to activate the Drosophila Toll signaling pathway (Buchon et al. 2014). If evolution of defense against *B. bassiana* was primarily through this humoral immune response, we may expect populations with increased defense against fungi to also have improved defense against Gram-positive *Enterococcus faecalis*, but we did not observe this. Evolved defense against fungus had no effect on defense against *E. faecalis*, nor against the Gram-negative bacterium *Providencia rettgeri*. Evolved defense may therefore be through other mechanisms, potentially including cellular immunity, melanization, composition of the cuticle, or even behaviors, such as grooming. Unlike our study, Wang et al. 2017 found that in the Drosophila Genetics Reference Panel (DGRP), defense against the fungus *Metarhizium anisopliae* Ma549 was positively correlated with defense against the Gram-negative bacterium *Pseudomonas aeruginosa* Pa14. Yet no correlation was seen between resistance to enteric infection with *P. entomophila* and inoculation (by stabbing) with *Erwinia carotovora* (Sleiman et al. 2015). It appears that *D. melanogaster* adaptation to parasites depends in part on the infection route, such that selection by oral infection against *Pseudomonas entomophila* did not confer resistance against systemic infection and vice versa (Martins et al. 2013). In our study, the two fungi were introduced by spray onto the fly cuticle. After contact with the cuticle, the fungus germinates and the hyphae penetrate the cuticle, presumably at multiple locations on the cuticle, and grow in the hemocoel of the fly. Our bacterial infections were done by pin prick into the fly thorax, thus leading to a localized wound on the cuticle. These different infection routes may be another reason for why we see no cross-resistance of fungal resistant populations against bacterial pathogens. But even with similar pathogens and similar infection routes, the same *D. melano*gaster genotypes are not resistant to all bacterial pathogens (Lazzaro et al. 2006). Defense against pathogens likely involves many genes, and potentially some of these may confer pathogen-specific defense, while others may contribute to some general aspect of robustness.

While several of the genes involved in *D. melanogaster* defense against fungal infection are known (Buchon et al. 2014), the genetic determinants of natural variation in a trait cannot be ascertained with experimental genetic disruptions. Experimentally induced mutations or gene knockdowns reveal extreme phenotypes due to loss of gene function but may not detect the complexity of gene regulation and interactions. From an evolutionary perspective, only a small subset of genes and processes that can be mutated to give a phenotype in the laboratory are expected to be responsible for segregating phenotypic variation in nature (Gruber et al. 2007). Furthermore, phenotypic variation can be shaped by genes that are not classically considered part of the immune system. Polymorphisms with smaller effect on immune defense may be identifiable in experimental evolution and resequencing studies, which can address the molecular architecture of adaptation. Previous experimental evolution and resequencing studies have not always implicated canonical genes that can affect a trait. When diverse base populations are used in thoughtfully-designed selection experiments (recommendations for experimental design are offered by Kofler and Schlotterer 2014 and Schlotterer et al. 2015), if known loci that affect a trait are not implicated, those loci are perhaps unlikely to be commonly involved in the trait in outbred populations where the loci are affected by forces of natural selection.

To identify candidate genes for immune defense, we compared SNP frequencies between our selected and control populations using CMH tests (Vlachos et al. 2019). We identified 45 significantly differentiated sites, even when using a simulation-based approach to correct for multiple comparisons, which resulted in a stringent significance threshold. Analyzing the significantly differentiated sites for GO term enrichment, while discarding GO terms with low representation and correcting for hierarchical clustering, we identified 29 enriched GO terms. These GO terms had a disproportionate involvement in membrane function. GO term analysis as applied to E&R studies such as ours implicitly assumes that many functionally related genes will be selected in the same way. But this assumption would only be true if the trait was determined by many functionally related genes that each contribute small effects to the adaptive phenotype. We do not necessarily expect this genetic architecture, and we consider this GO term analysis with some caution. If evolution of the trait is driven by a few genes with large allelic effects, then we might expect a lack of conspicuous GO enrichment, as we observe in our analysis.

It is worth noting that in this study we used larger population sizes than those commonly used in *D. melanogaster* experimental evolution studies, starting our selection protocol with 10,000 individuals in each replicate population. With large population sizes, four replicates per treatment, and nineteen generations of selection, we identified few candidate genes involved in immune defense against *B. bassiana*, despite the large phenotypic divergence between selected and control populations. This may be due to our study being underpowered by only having four replicate control populations compared with four replicate selected populations. While this level of replication is typical of existing E&R studies in *D. melanogaster*, some studies suggest that it may not be sufficient for detecting causal variants (Baldwin-Brown et al. 2014, Kofler and Schlotterer 2014). With current genomic tools, many more generations, and more replicates, may be needed to provide sufficient power to detect the many small-effect genes that are expected to confer immune defense.

Artificial selection experiments often display changes in SNP frequencies that appear to plateau before the end of the experiment, even as the phenotype continues to respond to selection. Parts et al. 2011 observed plateaus at intermediate allele frequencies in their yeast experimental evolution study, suggesting a reduction in selection coefficients. Such reductions in selection coefficients have been modeled by Illingworth et al. 2011, and plateaus in allele frequencies were also observed in *D. melanogaster* laboratory adaptation (Orozco-terwengel et al. 2012). We observed 4,000 sites across the entire genome that became fixed in the selected populations but were polymorphic in the ancestral and control populations. In comparison, only 1,000 sites fixed in the control populations but remained polymorphic in the ancestral and selected populations. Given the large effective population sizes in the S and C populations in every generation (N>1,000), the list of fixed sites in the S populations likely resulted from selection instead of drift, and may serve as candidate sites for immune defense variation.

The speed with which the selected populations evolved improved immune defense suggests the base population harbored ample genetic diversity. We might expect that other *D. melanogaster* populations may also harbor genetic diversity for immune defense. Among 297 Recombinant Inbred Lines from the Drosophila Synthetic Population Resource, we found vast levels of genetic diversity for 10-day survival post inoculation with the fungus *Beauveria bassiana*, ranging from 0 flies surviving to 100% of flies surviving the infection. Our QTL Mapping approach identified important genetic regions: one peak at a 5% FDR and three peaks at a 50% FDR. Under the 50% FDR, the protein coding genes included the Bomanin 55C gene cluster, specifically Bom1, Bom2, Bom3, Bom18 and Bom23. Expression of this gene cluster has been shown to be induced by *B. bassiana* infection (De Gregorio et al. 2001). Deletion of this cluster results in susceptibility to infection with fungi, although effects on resistance to *B. bassiana* were not directly tested (Clemmons et al. 2015, Lindsay et al. 2018). Immune deficiency (IMD) was also under the 50% FDR, along with odorant receptors (Or) and odorant binding proteins (Obp) which assist in the sensory perception of smell (Rollman et al., 2010). We also identified genes involved in the detection of taste, including Gustatory receptor (*Gr*), pickpocket (*ppk*) and ionotropic receptor (*Ir*) families, and genes involved in locomotion, including Ion transport peptide (ITP) (Hermann-Luibl, 2014), and TBPH (Feiguin 2009, Diaper et al., 2013). Under the 5% FDR were Jabba, which is thought to be involved in bacterial defense, genes involved in fatty acid elongation (CG18609 and CG17821), genes involved in transmembrane transporter activity (*MFS15*, *MFS14*, and *CG15096*) and an olfactory-expressed gene (*Mctp*). Since the RILs were lab adapted, it is possible that some immune defense variants from the wild were lost due to selection in the lab since immune defense likely comes at costs to other fitness characters. Nevertheless, ample variation for immune defense remained in the RILs.

Comparing the genomic regions identified in our E&R and QTL Mapping approaches, there were no overlaps. While both approaches resulted in a few candidate genes that we can follow up on in future studies, the lists of candidate genes were unique. Given the complexity of the immune defense phenotype, it is likely to be highly multigenic, and genes involved in this phenotype are likely to be involved in gene by gene, and gene by environment interactions. Thus, the lack of overlap between our two gene lists is not an indication of a lack of replication of these genes. Instead, both gene lists can be considered for further studies.

In the E&R study and in the QTL Mapping study, we saw evidence of sexual dimorphism in immune defense. A better understanding of what leads to sexual dimorphism in immune defense may guide the use of *B. bassiana* in biological control efforts that may benefit from targeting female insects. Previous studies have suggested that *D. melanogaster* females are more susceptible to infection with *B. bassiana* infection compared to males (Taylor and Kimbrell 2007, Kubiak et al. 2017, Shahrestani et al. 2018). Using the same *B. bassiana* ARSEF 12460 pathogen, we previously found that female flies were more susceptible to infection than male flies in inbred fly lines (Shahrestani et al. 2018), and this direction of sexual dimorphism was maintained whether the flies were sprayed or inoculated with the fungus, suggesting that grooming and barrier defenses are not fully responsible for sexual dimorphism in defense.

In our E&R study, there was sexual dimorphism in immune defense against *B. bassiana* only in the rate of evolution of cross-resistance to GHA, which evolved faster in males than females, when populations were selected to resist ARSEF 12460. Otherwise there was no sexual dimorphism in immune defense. In our QTL Mapping study, looking at 10-day post infection survival of the RILs, we found that the presence and direction of sexual dimorphism in immune defense is dependent on fly genotype. In approximately 50% of the RILs, females were more susceptible to infection than males, but in ∼25% of the RILs the pattern was reversed, and in the other ∼25% of the RILs there was no sexual dimorphism. This suggests that the direction of sexual dimorphism is not dependent just on a large-effect gene such as *relish*. The host genetic factors that affect the direction and magnitude of sexual dimorphism in immune defense remain a topic to investigate. One common hypothesis for sexual dimorphism in immune defense is differential reproductive investment of males and females leading to different resource allocation (reviewed in Schwenke et al. 2016). We do not have data for the reproductive output of the RILs used in this study, but it would be interesting to compare fecundity with sexual dimorphism in immune defense. Given the variation in both direction and magnitude of the observed sexual dimorphism, it is possible that several mechanisms can be involved in this trait. Despite the sexual dimorphism in defense, there was a positive correlation in 10-day survival of infection of males and females among the RILs.

Overall, we have shown that immune defense against a fungal pathogen is highly variable in *D. melanogaster* derived from natural populations. This variability allows rapid adaption in response to experimental selection, albeit at a cost to uninfected longevity. The presence of such extensive naturally occurring genetic variation suggests considerable adaptive potential in nature, although perhaps buffered by costs and tradeoffs. Notably, the variation appears to be distributed among multiple genes with modest allelic effects and no clear enrichment of functional gene categories. Nevertheless, using two different experimental approaches, we have identified a set of potentially causal genes that may be promising candidates for future study.

## Author Contributions

PS, BPL, and AGC conceived of the experiment. PS, KG, PC, GM, YE, MZ, SR, and MG performed the experiments and collected all data. EK, RR, MP, ZG, MR, and PS analyzed the data. PS, MP, MT, MR, AGC, and BPL wrote the manuscript. BPL, AGC, JV, SW, and ADL provided training, expertise, and resources.

## Acknowledgements

This project was funded by NIH F32 GM109700 (PS and BPL) and R01 AI083932 (BPL). The following students from the Shahrestani lab contributed to various aspects of the project, including presenting results at local conferences: Andrew Talbott, Christian Gochez, Julianne Kim, Hannah Ro, and Julie Nguyen.

---

**Figure S1.**
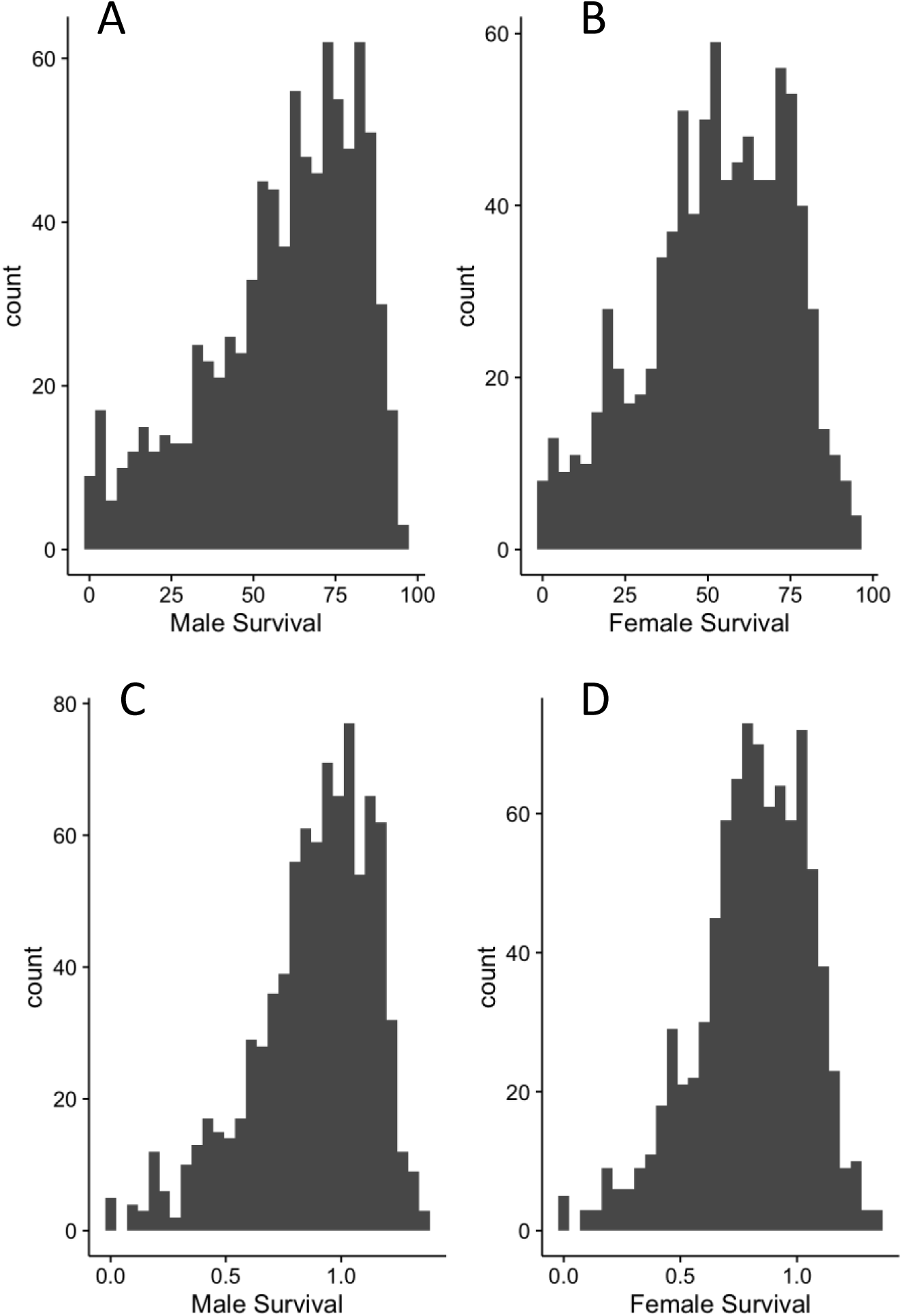
Distribution of 10th-day survival of the DSPR RILs post-inoculation with *B. bassiana* ARSEF 12460. The distribution was slightly skewed for both males (A) and females (B), as seen with the count of RILs plotted against the survival percentage. We used an arcsin square root transformation of survival percentage for both males (C) and females (D).

**Figure S2.**
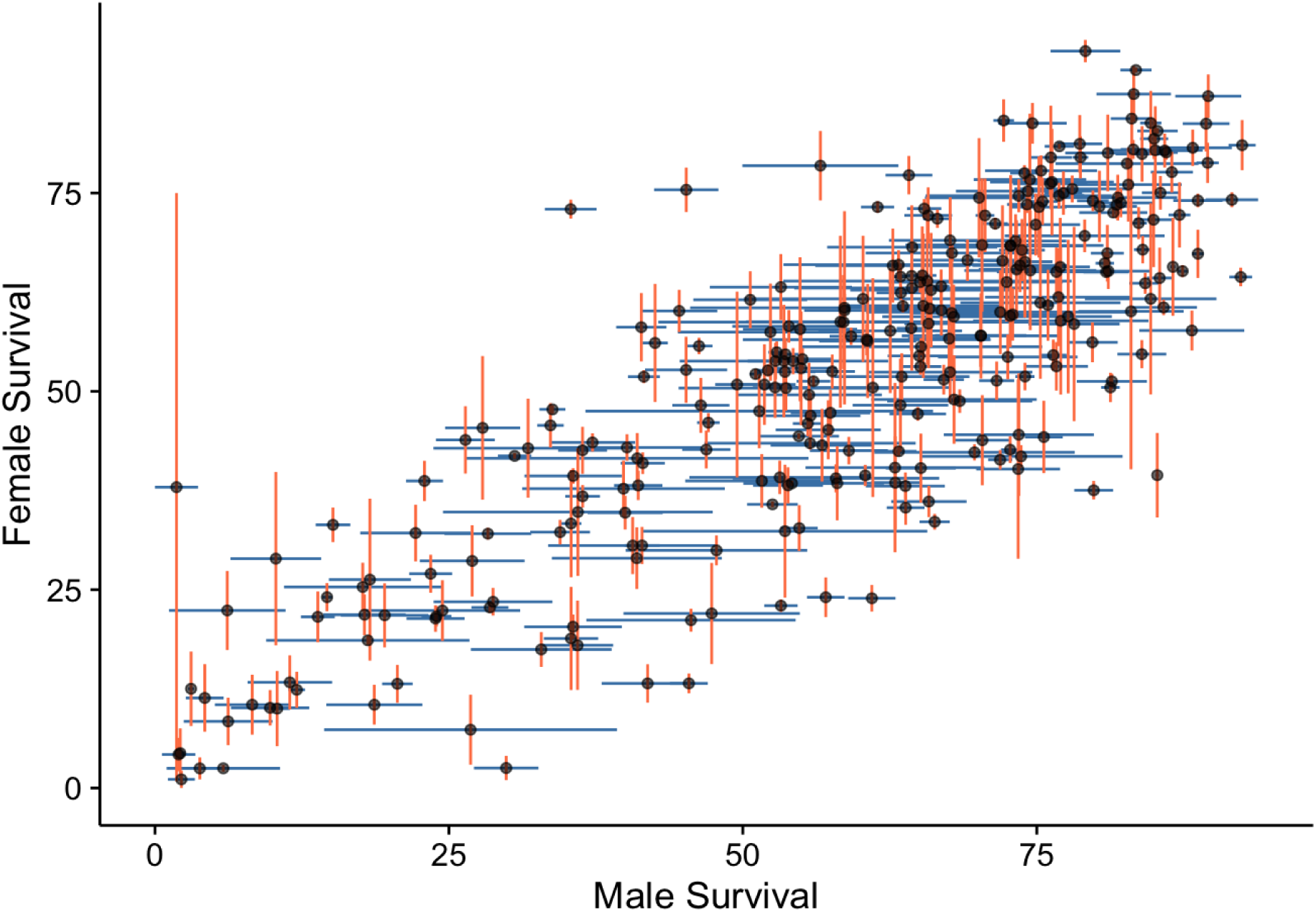
Correlation between male (blue) and female (red) survival 10 days after inoculation with *B. bassiana* ARSEF 12460 (*r*^2^ = 0.59; p < 2.2×10^-16^). Lines show the standard error of the mean for females (vertical) and males (horizontal).

**Figure S3.**
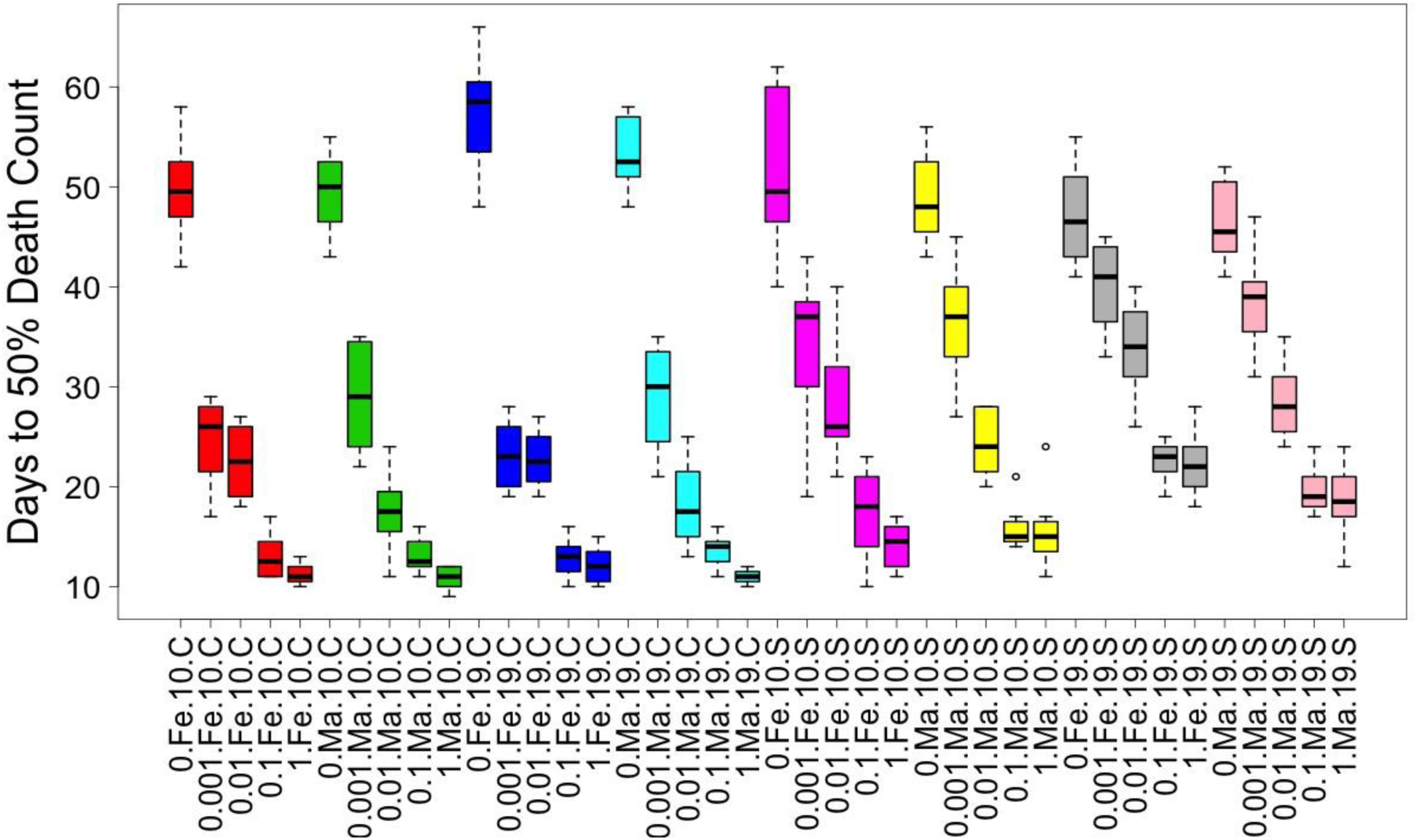
LT_50_ categorized by population (C_1-4_ vs. S_1-4_), generation (10 vs. 19), sex, and dose. Within each color, the 5 boxplots represent the 5 doses as proportions relative to the highest dose of 10^4^ spores/mm^2^: 0, 0.001, 0.01, and 1. The left half of the X axis shows the C populations and the right half shows the S populations. Within each population type, the first half are generation 10 data and the second half are generation 19 data. After 19 generations of selection, both males and females from the selected (S) populations became more robust to every dose. When uninfected (dose 0), the S populations had lower LT50 compared to the C populations after 19 generations of selection.

**Figure S4.**
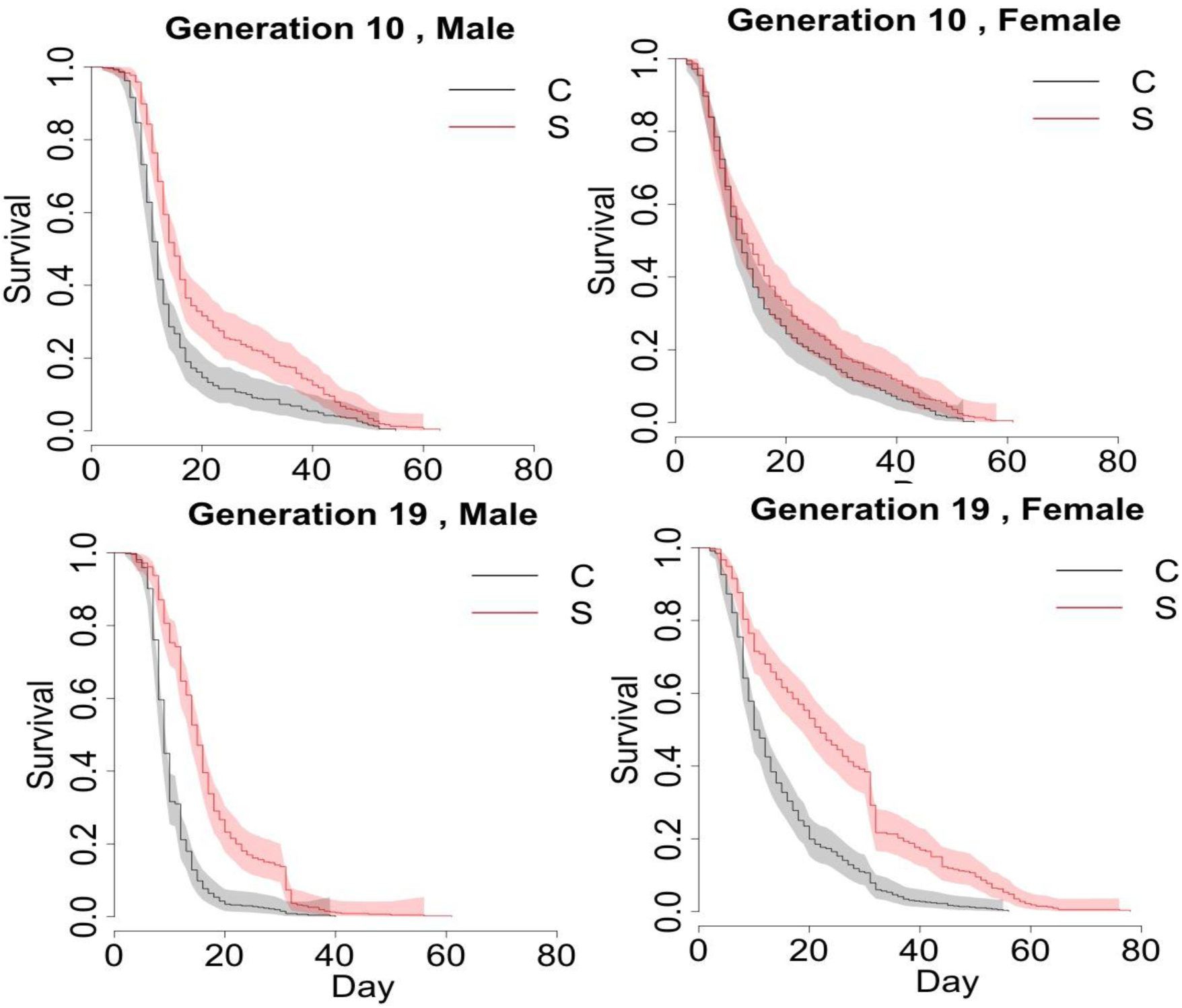
The Kaplan-Meier estimate of the survival function is shown for S_1-4_ (red) and C_1-4_ (black) populations across different combinations of sex (columns) and generation (rows). After 19 generations of selection for defense against *B. bassiana* ARSEF 12460, both males and females evolved cross-resistance against *B. bassiana* GHA. However, the rate of this evolution differed between males and females (Table 3, sex by generation interaction: p<0.0001)

**Figure S5.**
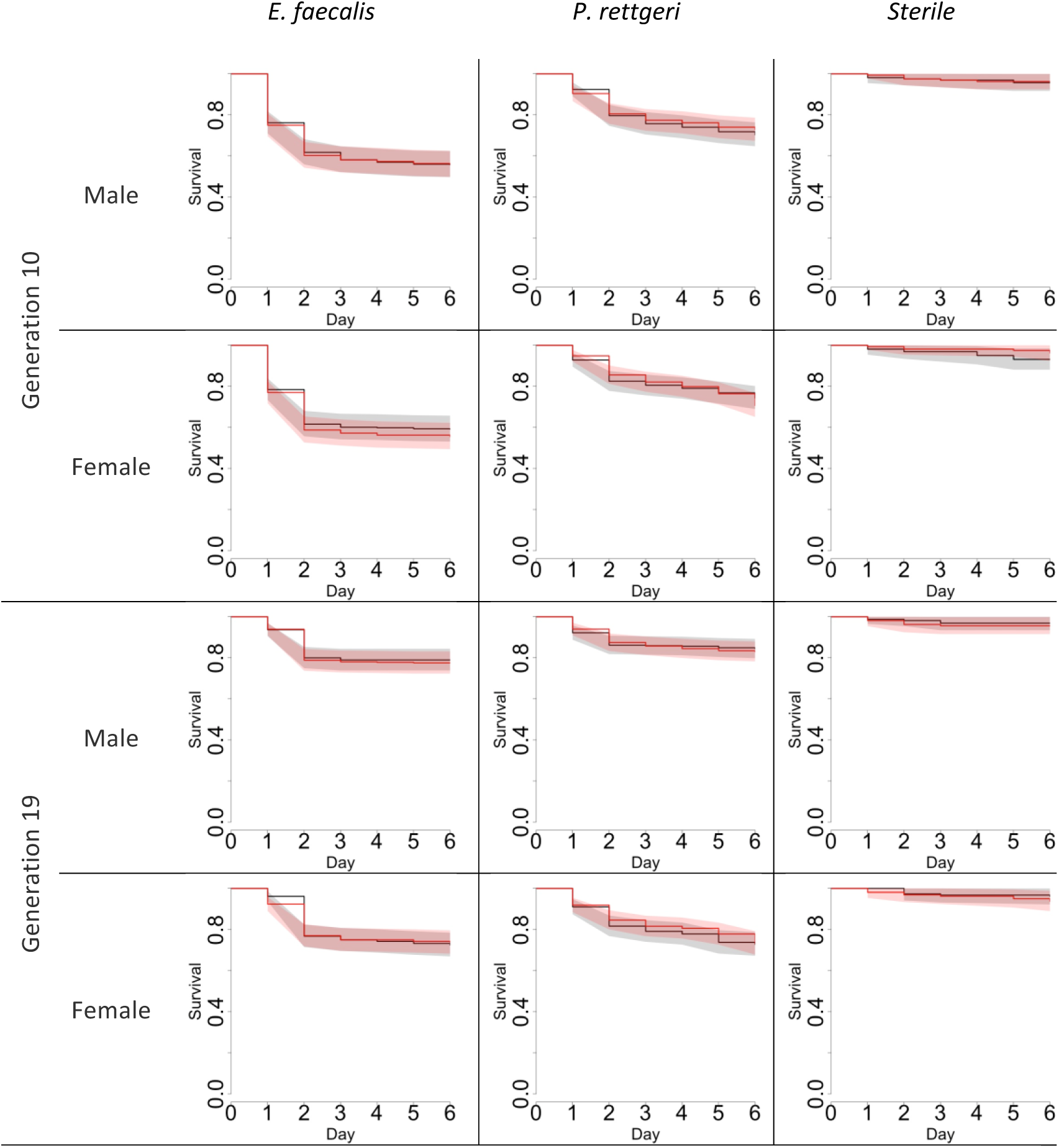
The Kaplan-Meier estimate of the survival function for S and C populations across different combinations of sex, generation, and infection: *E. faecalis* (left), *P. rettgeri* (center), and uninfected sterile (right). The shaded areas show 95% confidence intervals. When compared to the sterile group, the two bacterial infected groups die faster (p<0.0001 from log-rank test for both generations 10 and 19).

**Figure S6.**
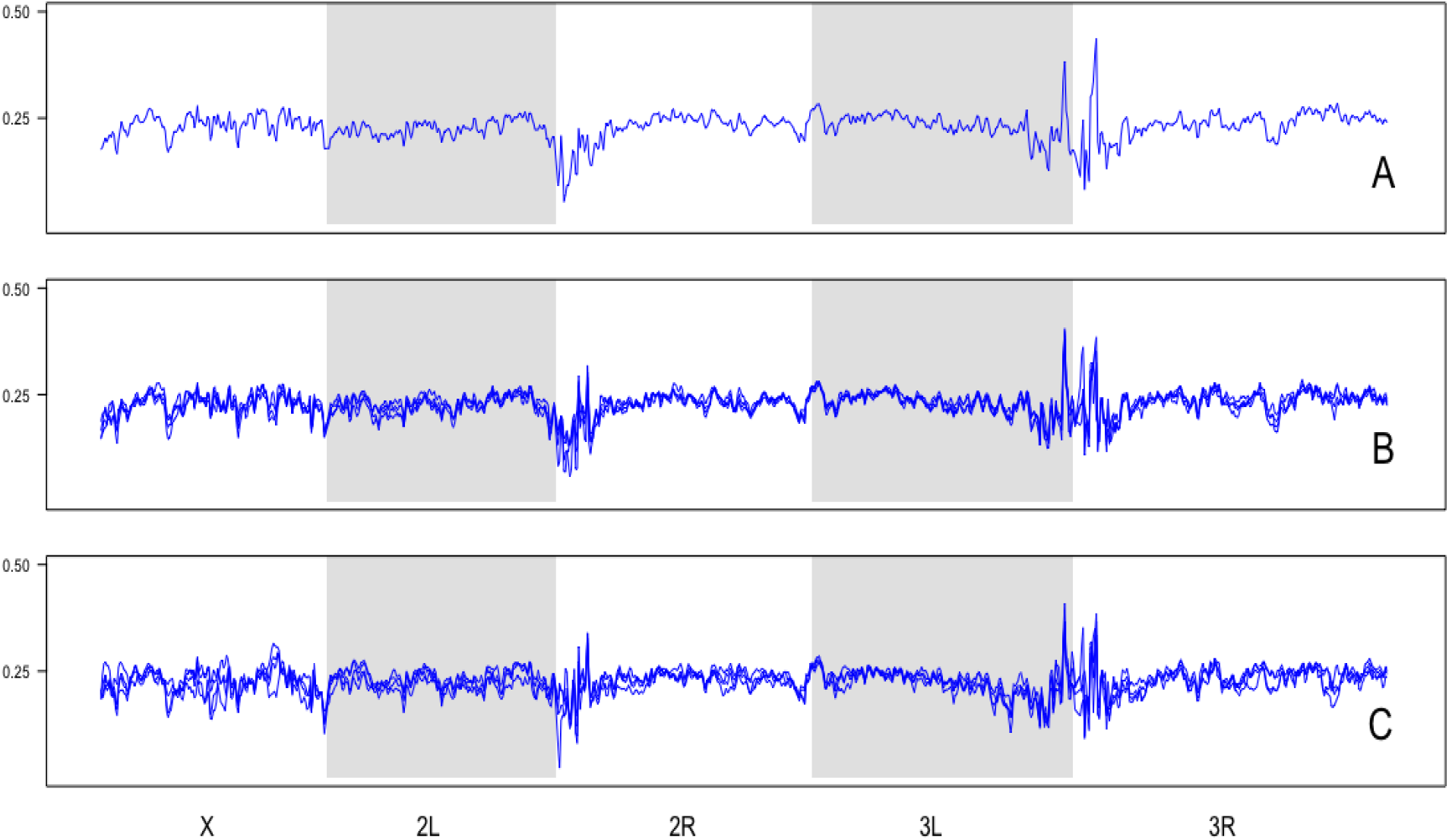
Heterozygosity in the ancestral P *(A)*, control C *(B)*, and selected S (*C)* populations plotted over 150-kb windows across all major chromosome arms. All replicates are shown for each population.

**Figure S7.**
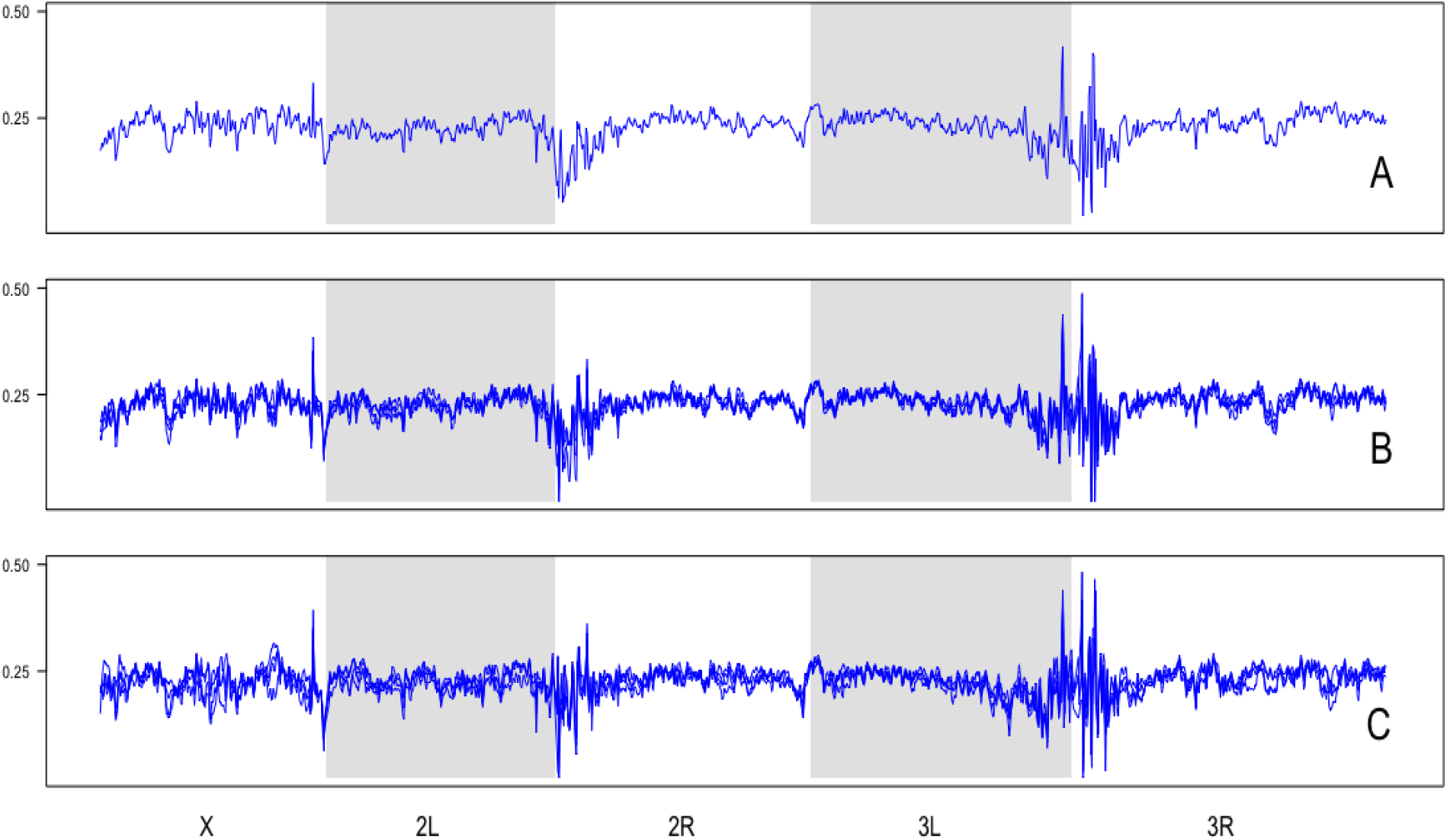
Heterozygosity in the ancestral P *(A)*, control C *(B)*, and selected S (*C)* populations plotted over 100-kb windows across all major chromosome arms. All replicates are shown for each population.

**Figure S8.**
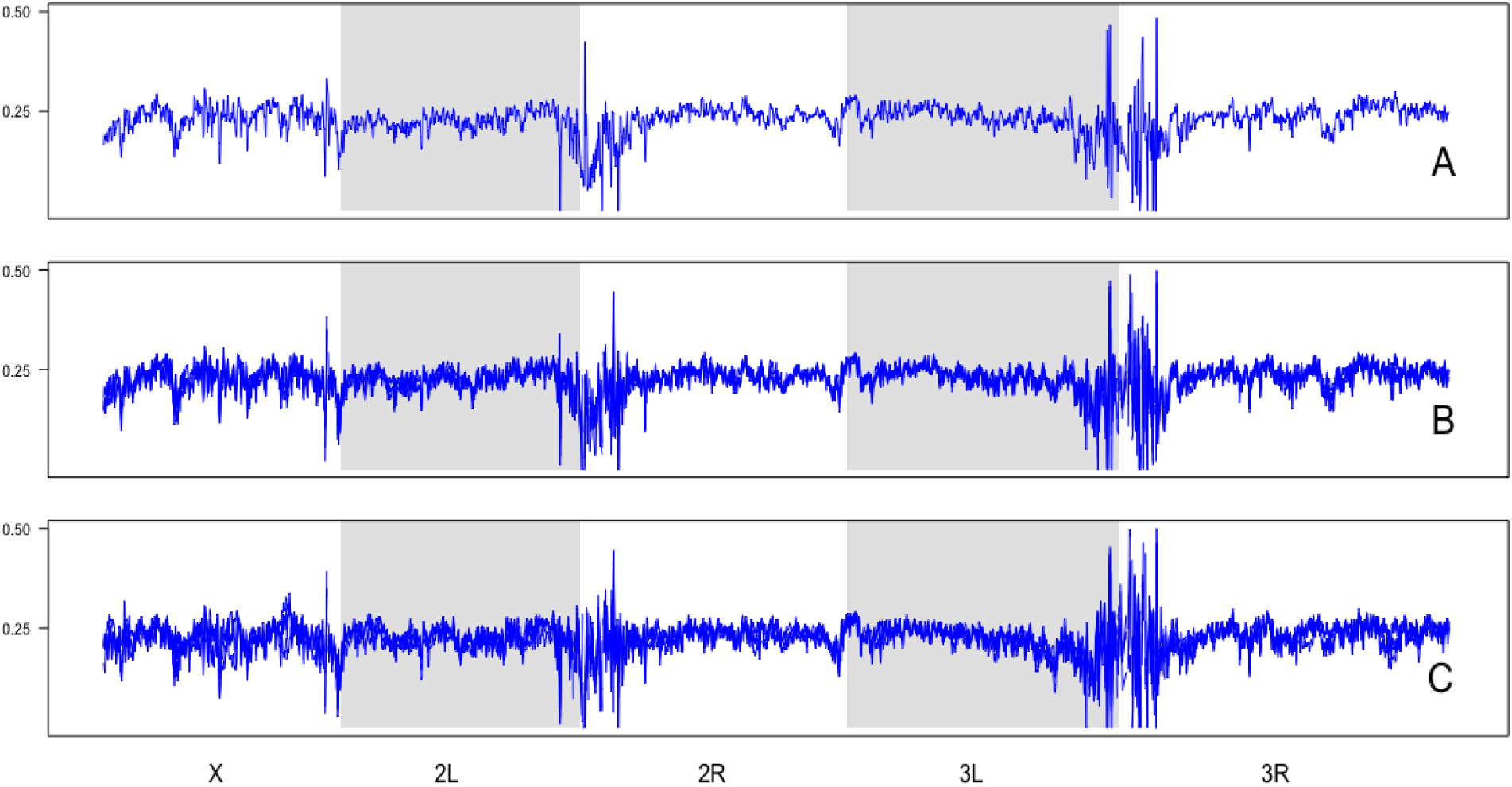
Heterozygosity in the ancestral P *(A)*, control C *(B)*, and selected S (*C)* populations plotted over 50-kb windows across all major chromosome arms. All replicates are shown for each population.

**Figure S9.**
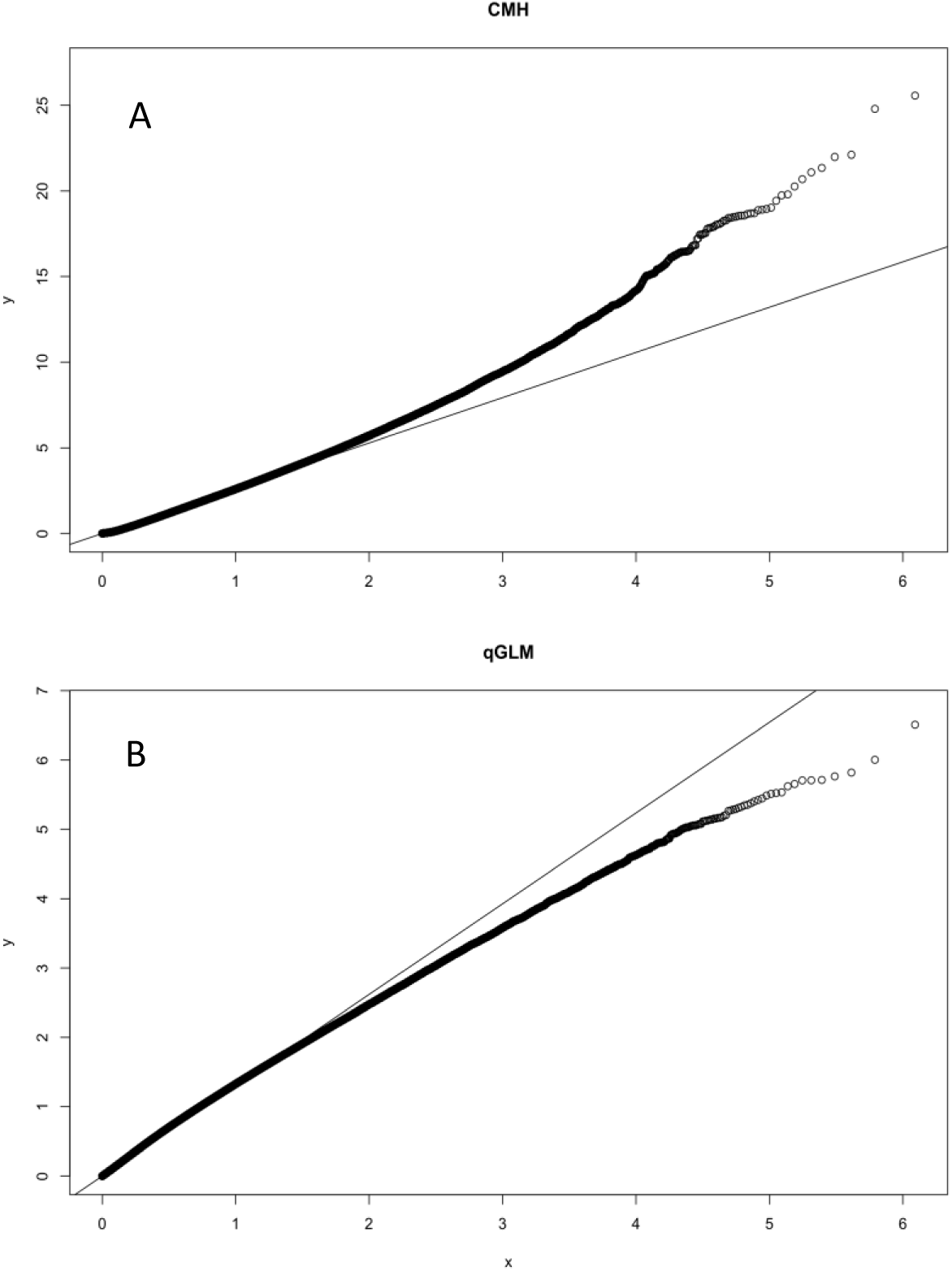
Q-Q plots of observed vs. expected p values in the CMH analysis (A) and qGLM analysis (B). Q-Q plots suggest that the CMH test is better suited for our data than the qGLM method. Note the difference in the y axes.

**Table S1.** Average read coverage across the genome for the C, S, and P populations, and mean genome-wide heterozygosity based on SNP frequencies.

| **Population** | **Replicate** | **Average Read Coverage** | **Mean Heterozygosity** |
| --- | --- | --- | --- |
| C | 1 | 45 | 0.24 |
|  | 2 | 44 | 0.24 |
|  | 3 | 25 | 0.23 |
|  | 4 | 35 | 0.23 |
| S | 1 | 45 | 0.23 |
|  | 2 | 59 | 0.24 |
|  | 3 | 34 | 0.23 |
|  | 4 | 32 | 0.23 |
| P | 1 | 36 | 0.24 |

**Table S2.** Coordinates of QTL peaks. The QTL Mapping was done separately for males, females, and the difference between males and females (dimorphism). At 50% FDR, one significant peak was identified in each group. At the 5% FDR, one significant peak was identified in the males. The analysis output 5.x coordinates, which were converted to 6.x coordinates in Flybase prior to searching for genes in these regions.

| **Sex** | **FDR** | **5x coordinates (result)** | **6x conversion** |
| --- | --- | --- | --- |
| Male | 5% | 2R:14,560,000..14,700,000 | 2R:18,672,495..18,812,495 |
| Male | 50% | 2R:15,740,000..16,590,000 | 2R:19,852,495..20,702,495 |
| Female | 50% | 2R:14,120,000..14,410,000 | 2R:18,232,495..18,522,495 |
| Dimorphism | 50% | 2R:19,480,000..20,420,000 | 2R:23,592,477..24,532,477 |

Table S3: See Supplemental file 1

**Table S4.**
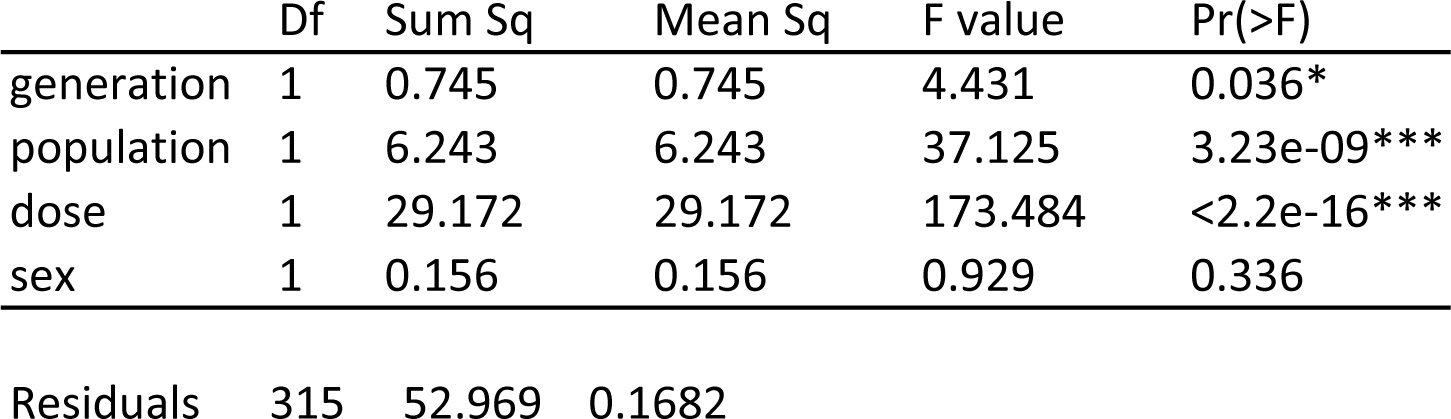
ANOVA model testing the effects of sex, generation, population, and dose of *B. bassiana* ARSEF 12460 on LT50.

**Table S5.**
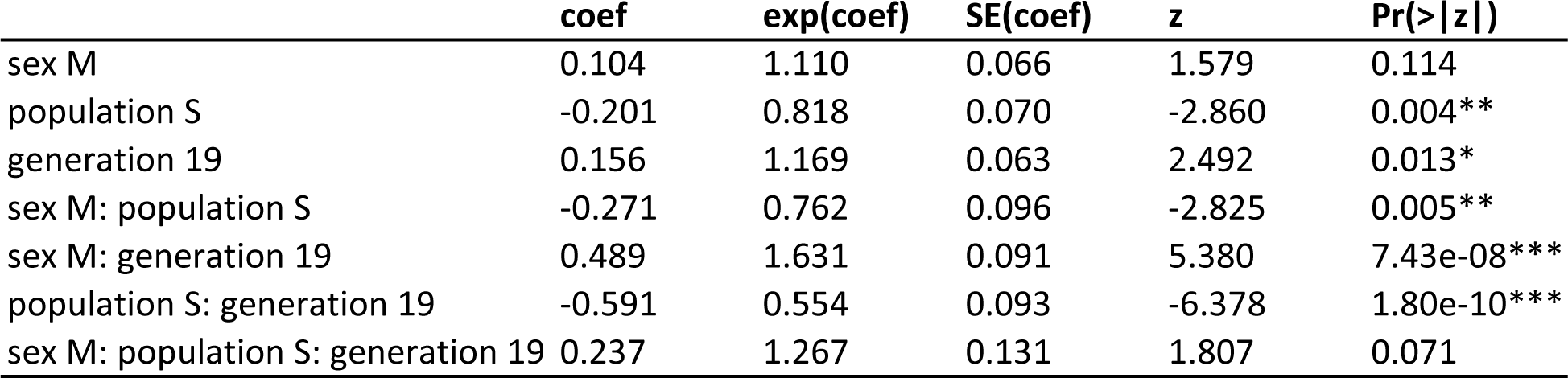
Analysis of survival differences in selected and control populations with and without infection at generations 10 and 19 with and without infection with B. bassiana GHA. Survival was analyzed using model 2: coxph = S + P + G + S*P + S*G + P*G+ S*P*G, where G represents generation (10 and 19), P represents populations (C and S), and D represents the infection status (uninfected and infection with 104 spores/mm2). n= 3849.

|  | coef | exp(coef) | SE(coef) | z | Pr(>\|z\|) |
| --- | --- | --- | --- | --- | --- |
| sex M | 0.104 | 1.110 | 0.066 | 1.579 | 0.114 |
| population S | -0.201 | 0.818 | 0.070 | -2.860 | 0.004** |
| generation 19 | 0.156 | 1.169 | 0.063 | 2.492 | 0.013* |
| sex M: population S | -0.271 | 0.762 | 0.096 | -2.825 | 0.005** |
| sex M: generation 19 | 0.489 | 1.631 | 0.091 | 5.380 | 7.43e-08*** |
| population S: generation 19 | -0.591 | 0.554 | 0.093 | -6.378 | 1.80e-10*** |
| sex M: population S: generation 19 | 0.237 | 1.267 | 0.131 | 1.807 | 0.071 |

**Table S6.** Hazard ratios between the S and C populations across different groups formed by sex and generation, for infection with GHA.

|  | Generation 10 |  | Generation 19 |  |
| --- | --- | --- | --- | --- |
|  | Male | Female | Male | Female |
| Hazard Ratio (S vs. C) | 0.62 | 0.82 | 0.44 | 0.45 |
| p-value | 0.0002 | 0.0243 | <0.0001 | <0.0001 |

Table S7. See Supplemental file 1

Table S8. See Supplemental file 1

Table S8. See Supplemental file 2

**Table S10.** Enriched GO terms from genomic differentiation of S vs. C populations

| GO ID | P-value | GO Term |
| --- | --- | --- |
| GO:0019725 | 0.000849 | cellular homeostasis |
| GO:0009306 | 0.001393 | protein secretion |
| GO:0090175 | 0.002742 | regulation of establishment of planar polarity |
| GO:0007160 | 0.003197 | cell-matrix adhesion |
| GO:0051130 | 0.006269 | positive regulation of cellular component organization |
| GO:0098590 | 0.006325 | plasma membrane region |
| GO:0098589 | 0.006512 | membrane region |
| GO:0043604 | 0.006637 | amide biosynthetic process |
| GO:0044089 | 0.00881 | positive regulation of cellular component biogenesis |
| GO:0051649 | 0.011888 | establishment of localization in cell |
| GO:0045887 | 0.013467 | positive regulation of synaptic growth at neuromuscular junction |
| GO:1904398 | 0.013467 | positive regulation of neuromuscular junction development |
| GO:0051965 | 0.014311 | positive regulation of synapse assembly |
| GO:0016323 | 0.014828 | basolateral plasma membrane |
| GO:0097060 | 0.019744 | synaptic membrane |
| GO:0051124 | 0.020918 | synaptic growth at neuromuscular junction |
| GO:0012505 | 0.021673 | endomembrane system |
| GO:0007617 | 0.02192 | mating behavior |
| GO:0042626 | 0.027998 | ATPase activity, coupled to transmembrane movement of substances |
| GO:0003824 | 0.031367 | catalytic activity |
| GO:0016324 | 0.032655 | apical plasma membrane |
| GO:0016887 | 0.032792 | ATPase activity |
| GO:0044444 | 0.033109 | cytoplasmic part |
| GO:0051094 | 0.037168 | positive regulation of developmental process |
| GO:0008049 | 0.039698 | male courtship behavior |
| GO:0048471 | 0.040054 | perinuclear region of cytoplasm |
| GO:0097458 | 0.042283 | neuron part |
| GO:0022603 | 0.045684 | regulation of anatomical structure morphogenesis |
| GO:0044802 | 0.046314 | single-organism membrane organization |

**Table S11.**
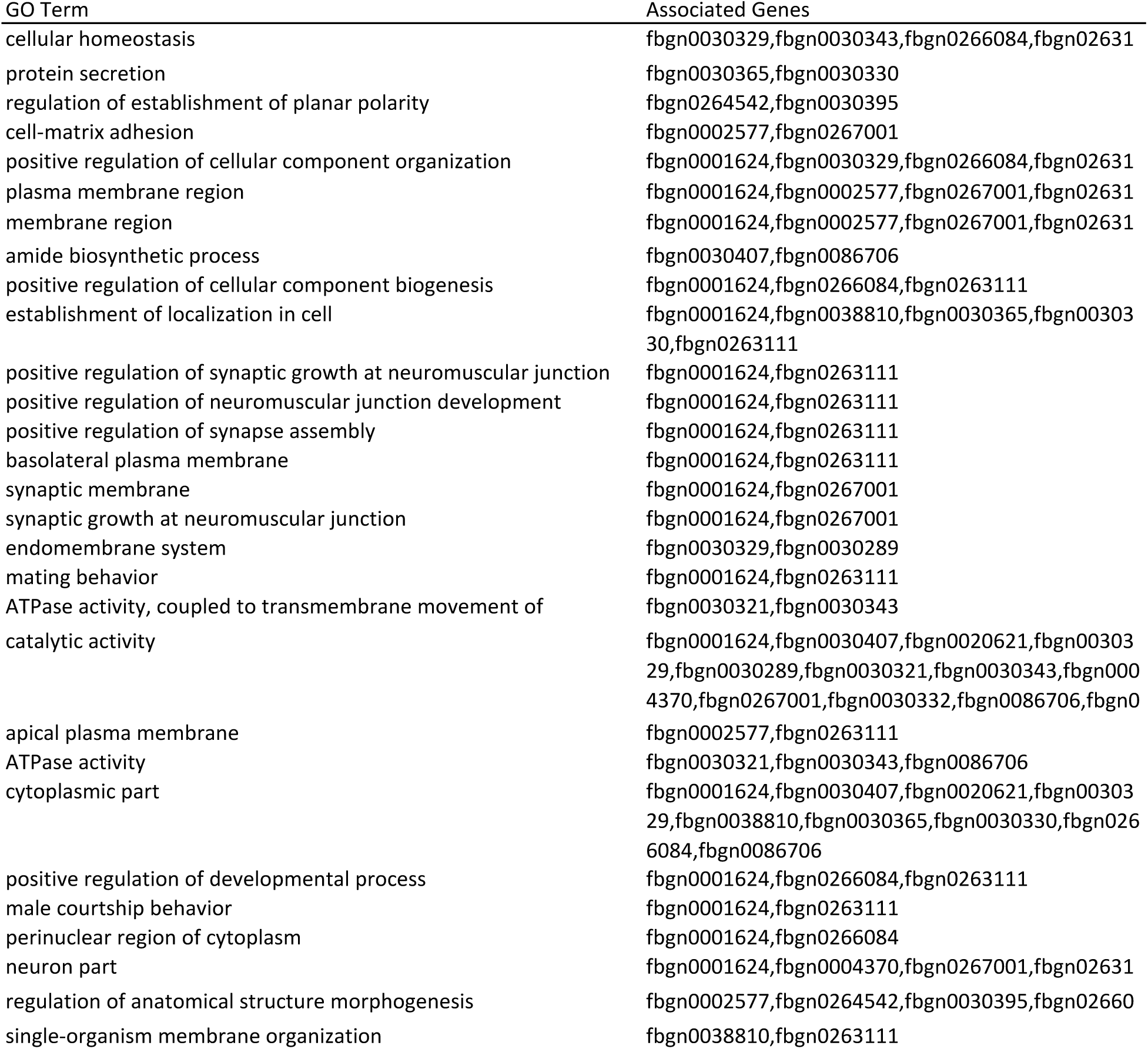
Genes associated with enriched GO terms from S vs. C population comparison

